# Allosteric communications between domains modulate the activity of matrix metalloprotease-1

**DOI:** 10.1101/804559

**Authors:** Lokender Kumar, Anthony Nash, Chase Harms, Joan Planas-Iglesias, Derek Wright, Judith Klein-Seetharaman, Susanta K. Sarkar

## Abstract

An understanding of the structure-dynamics relationship is essential for understanding how a protein works. Prior research has shown that the activity of a protein correlates with intra-domain dynamics occurring at picosecond to millisecond timescales. However, the correlation between inter-domain dynamics and the function of a protein is poorly understood. Here we show that communications between the catalytic and hemopexin domains of matrix metalloprotease-1 (MMP1) on type-1 collagen fibrils correlate with its activity. Using single-molecule FRET (smFRET), we identified functionally relevant open conformations where the two MMP1 domains are well-separated, which were significantly absent for catalytically inactive point mutant (E219Q) of MMP1 and could be modulated by an inhibitor or an enhancer of activity. The observed relevance of open conformations resolves the debate about the roles of open and closed MMP1 structures in function. A sum of two Gaussians fitted histograms, whereas an exponential fitted autocorrelations of smFRET values. We used a two-state Poisson process to describe the dynamics and used histograms and autocorrelations of conformations to calculate the kinetic rates between the two states. All-atom and coarse-grained simulations reproduced some of the experimental features and revealed substrate-dependent MMP1 dynamics. Our results suggest that an inter-domain separation facilitates opening up the catalytic pocket so that the collagen chains come closer to the MMP1 active site. Coordination of functional conformations at different parts of MMP1 occurs via allosteric communications that can take place via interactions mediated by collagen even if the linker between the domains is absent. Modeling dynamics as a Poisson process enables connecting the picosecond timescales of molecular dynamics simulations with the millisecond timescales of single molecule measurements. Water-soluble MMP1 interacting with water-insoluble collagen fibrils poses challenges for biochemical studies that the single molecule tracking can overcome for other insoluble substrates. Inter-domain communications are likely important for multidomain proteins.

**Statement of Significance:** It is often challenging to distinguish functionally important dynamics because proteins are inherently flexible. MMP1 is a model enzyme because both the catalytic and hemopexin domains are necessary to degrade triple-helical type-1 collagen, the highly proteolysis-resistant structural component of the extracellular matrix. We report, for the first time, measurements of MMP1 inter-domain dynamics on type-1 collagen fibrils. We have identified functionally relevant MMP1 conformations where the two domains are far apart. Mutations and ligands can allosterically modulate the dynamics that correlate with activity. The dynamics follow a two-state Poisson process that connects the picosecond timescales of MD simulations with the millisecond timescales of experiments. The two domains can functionally communicate via collagen even when the physical linker is absent.

## Introduction

Researchers have argued both in favor of (1) and against (2) the roles of protein conformational dynamics in the activity. A direct link between the timescale hierarchies of intra-domain dynamics from pico- to milli-second and function has been argued based on experimental and computational studies (3). However, researchers do not entirely know how the inter-domain dynamics of a protein at different timescales influence its activity. Collagen degradation by MMP1 provides a unique opportunity to define the relationship between inter-domain dynamics and function because both the catalytic and hemopexin domains of MMP1 are necessary to degrade structurally intact collagen (4). Collagen is the primary component of the extracellular matrix (ECM) that provides a scaffold for cells to maintain tissue integrity. Degradation of fibrils by matrix metalloproteases (MMPs) is an integral part of tissue remodeling (5). MMP1, a collagenase in the 23-member MMP family, can degrade type-1 collagen. MMP1 consists of a catalytic domain that degrades collagen, a hemopexin domain that helps MMPs bind to collagen, and a linker that mediates communications between the two (6). Even though MMPs have differences in activity and substrate specificity, the catalytic domain sequence across the MMP family is very similar to MMP1 (7). The functional difference, despite the catalytic domain similarity, suggests that the two domains communicate allosterically, which means a signal originating at one site transfers to a distant site where the protein’s function such as catalysis occurs (8).

Most studies on MMP1-collagen interactions have used water-soluble collagen monomers instead of water-insoluble collagen fibrils. Each 300 nm long collagen monomer of diameter 1.5 nm consists of three left-handed chains forming a right-handed triple-helical structure that hides the cleavage sites and makes collagen resistant to degradation (9). Biochemical studies on collagen monomers revealed that MMP1 actively unwinds the three strands (4). Follow-up studies reinterpreted the same results (4) to introduce the concept of collagen breathing, which leads to a partial unwinding of collagen due to thermal fluctuations, enabling MMP1 to bind and cleave collagen (10). Additional experimental studies supported such passive unwinding of collagen due to temperature (11). Although the catalytic domain alone can degrade denatured collagen, it can not degrade triple-helical collagen without the hemopexin domain (4). The essential role of the hemopexin domain in function suggests communications between distant parts of MMPs. Further research on MMP1 (12, 13) and MMP3 (14) provided additional support for allosteric interactions between domains. One set of studies suggests that a larger separation (open conformation) between the catalytic and hemopexin domains is necessary for activity based on NMR and SAXS experiments and all-atom molecular dynamics (MD) simulations (13, 15-17). Another set of studies suggests that the two domains are closer (closed conformation) based on X-ray crystallography (12). We need to resolve the debate about the open and closed conformations.

However, the process of degrading physiologically relevant collagen fibrils is different from that of degrading monomers for several reasons. First, collagen fibrils are water-insoluble and macroscopic, greatly complicating ensemble biochemical and kinetics studies. Fibrils are larger than MMPs in size, and MMPs bind to fibrils nonspecifically as well as at the partially unwound vulnerable sites created during fibril assembly (18, 19). As such, binding to the cleavage sites is not a simple diffusion-limited process determined by the diffusion constant in solution. Also, the combined MMP1-fibril system is not statistically time-independent because the fibril itself changes upon degradation by MMP1. Second, as monomers self-assemble into fibrils, the cleavage sites become less accessible compared to monomers due to covering by the C-terminal telopeptides (20). Saffarian et al. (21) showed that collagen degradation biases the motion of MMP1 on type-1 collagen fibrils using Fluorescence Correlation Spectroscopy (FCS). Using single-molecule tracking of labeled MMP1 on type-1 collagen fibrils, we showed that MMP1 diffusion is both biased and hindered due to cleavage (19). MMP1 spends more than 90% of its time on the collagen fibril by diffusing, binding, and pausing without initiating degradation. Such extensive binding and pausing reduce the apparent catalytic rate of MMP1. The subsequent publication showed that fibrils also play a role in MMP1 activity due to vulnerable sites on fibrils caused by the relaxation of strain during fibril assembly (18). In summary, the overall catalytic rate of MMP1 on fibrils depends on random motion (19) and substrate properties (18) as well as on active site catalysis (12) and conformational dynamics (13) applicable to both fibrils and monomers. However, there is no report of MMP1 conformational dynamics on fibrils and their roles in the active site catalysis.

Here we focus on collagen fibrils as opposed to collagen monomers and show that functionally relevant MMP1 conformations have the catalytic and hemopexin domains distant. These conformations are present in active MMP1 but are significantly absent in inactive MMP1. Tetracycline, an antibiotic that is known to inhibit MMP1 activity (22, 23), inhibits these conformations. In contrast, MMP9, another member of the MMP family that cannot degrade triple-helical collagen (24), enhances the low FRET conformations. Molecular dynamics simulations of active and inactive MMP1 bound to a model triple-helical collagen reproduced some of the experimental observations. MD simulations further revealed that MMP1 opens its catalytic domain more compared to inactive MMP1. Also, the root-mean-squared (rms) distance between the MMP1 catalytic site and the cleavage sites on the collagen chains has a lower average value for active MMP1 compared to inactive MMP1.

## Materials and methods

### Selection of labeling sites on MMP1

For a fluorescence-based study of inter-domain motions, fluorescent dyes need to be attached to MMP1. For this purpose, we analyzed the crystal structure of MMP1 bound to a model of triple-helical collagen monomer (PDB ID 4AUO) for suitable sites for labeling on the catalytic and hemopexin domains. Since there are numerous surface-exposed lysine residues, labeling using dyes with NHS ester is not appropriate. Since disulfide bonds between cysteine residues are generally crucial for structural stability (25), two cysteine residues (CYS278 and CYS466) joined by a disulfide bond in MMP1 are not good candidates for mutations. However, CYS278 and CYS466 are relatively hidden, making them less labile. Consequently, we chose to mutate one serine residue from each domain to cysteine residues. We considered several factors before selecting the two sites for mutations: 1) the distance between the locations should be around the Förster radius of the dye pair chosen for smFRET, 2) the amino acid at the site should be solvent-exposed to facilitate labeling, 3) the amino acid selected should be similar to cysteine to minimize the effect of a mutation on activity, 4) the amino acid should not be in a conserved region, 5) the amino acid should not be near the collagen-binding site, and 6) the amino acid should preferably be on the relatively stable helices. Based on these criteria, we mutated SER142 on the catalytic domain and SER366 on the hemopexin domain to cysteines (**Fig. 1 A**).

**Figure 1.**
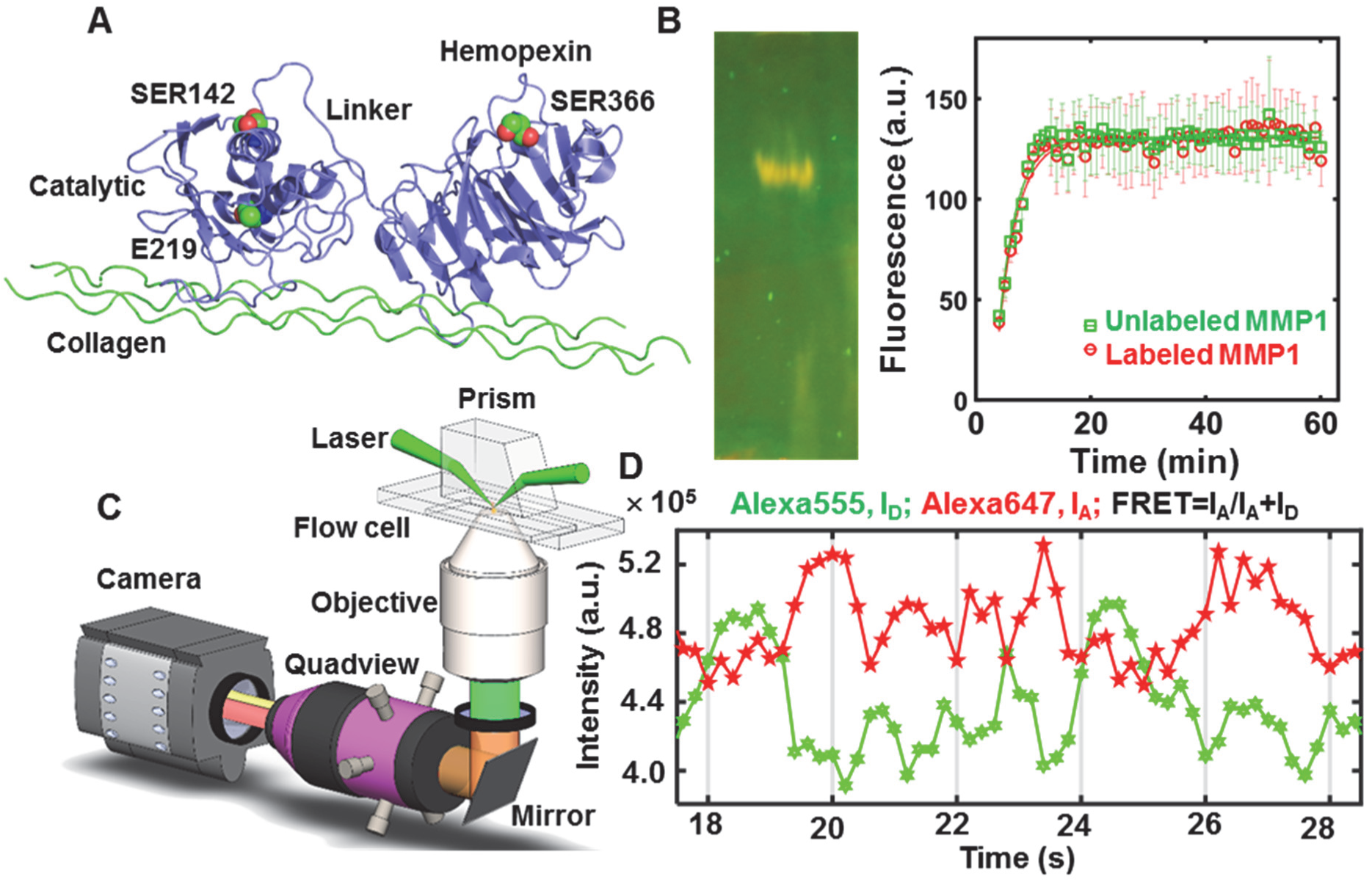
Single-molecule measurement of MMP1 dynamics on type-1 collagen fibril. (**A**) Relative positions of the MMP1 domains and residues (green and red spheres) created using 4AUO. (**B**) Left panel: 12% SDS PAGE of labeled MMP1; Right panel: Fluorescence from degraded peptide substrate as a function of time for unlabeled (green squares) and labeled (red circles) MMP1 at 37° C. Solid lines are respective best fits to *y=a-b*\*exp(-*kt*). After calibration, the specific activity is ∼1000 pmol/min/μg. The error bars are the stds of 3 technical repeats. (**C**) Schematics of the TIRF microscope. (**D**) Emission intensities of the two dyes attached to active MMP1. Low FRET conformations lead to high Alexa555 emission, whereas high FRET conformations lead to low Alexa555 emission. Anticorrelated Alexa647 and Alexa555 emissions, I_A_ and I_D_, respectively, indicate the conformational dynamics of MMP1.

### Purification, labeling, and ensemble activity measurements

We expressed full-length recombinant MMPs with pro domains in *E. coli*. We cleaved off pro domains using 0.1 mg/mL trypsin to activate both active and catalytically inactive point mutant (E219Q) MMP1 as well as active MMP9 (26). We purified activated MMPs using a protease based purification method (27). We labeled purified MMP1 with AlexaFluor555 (ThermoFisher Scientific, Cat# A20346, donor dye) and AlexaFluor647 (ThermoFisher Scientific, Cat# A20347, acceptor dye) using maleimide chemistry. 1 mL of MMP1 at a concentration of 1 mg/mL was incubated with 20 µL of 1 mg/mL AlexaFluor555 and AlexaFluor647 for 60 minutes in a 5 mL glass vial with continuous nitrogen flow to avoid oxidation of the dyes. After incubation, we filtered the sample three times using a 30 kDa cut-off Amicon filter to remove free dyes from the solution. We analyzed the integrity of labeled MMP1 using 12% SDS PAGE (**Fig. 1 B**, left panel). We compared the activities of labeled MMP1 and unlabeled MMP1 (**Fig. 1 B**, right panel) on the synthetic substrate, MCA-Lys-Pro-Leu-Gly-Leu-DPA-Ala-Arg-NH2, (R&D Systems, Cat# ES010) (27). MMP9 was not labeled.

### Single-molecule measurements

We mixed 90 µL of 4 mg/mL solution of type-1 collagen monomers (Advanced Biomatrix, RatCol type-1 collagen for 3D gels, Cat# 5153) with 10 µL of neutralizing solution (Advanced Biomatrix, Neutralization Solution, Cat# 5155) to create a reaction volume of 100 µL. We created a thin layer of reconstituted type-1 collagen fibrils on a quartz slide using the 100 µL reaction. After incubation for 1 h at 37° C, we made a flow cell using a piece of double-sided tape sandwiched between the quartz slide and a glass coverslip. We mixed 50 µL of 0.1 mg/mL labeled MMP1 with 50 µL of 1) protein buffer (50 mM Tris, 100 mM NaCl, pH 8.0), 2) 1 mg/mL MMP9, and 3) 100 µg/mL tetracycline. We incubated the mixtures for 30 min at 22° C. We serially diluted the labeled MMP1 to prepare a concentration of ∼100 pM. Before flowing in labeled MMPs into the flow cell, we photobleached the slide using the highest available power of the laser in our TIRF microscope (25 mW laser power at 532 nm focused using a 10 cm plano-convex lens). As a result, bright spots due to impurities photobleached and image frames became spot-free. Then, we flowed labeled MMPs and acquired movies. Alexa555 dyes attached to MMP1 were excited at 532 nm using the evanescent wave created at the interface of the quartz slide and sample solution in a Total Internal Reflection Fluorescence (TIRF) Microscope as described before (18, 19). We separated Alexa647 and Alexa555 emissions into different channels using Quad View Simultaneous Imaging System (Photometrics, QV2) and imaged using an EMCCD camera (Andor iXon) at 22° C with 100 ms time resolution. We superimposed the two emission channels using a pairwise stitching plugin of ImageJ and extracted the emission intensities from the two dyes for further analysis using Matlab. For each experimental condition (no ligand, with MMP9, and with tetracycline), we acquired at least 10 movies with 100 ms time resolution with 3000 frames for both active and inactive MMP1. The number of anticorrelated trajectories was ∼20±14% (mean±standard deviation (std)) of the total number of spots in a movie.

### Analysis of potential labeling issues

Upon activation by trypsin, MMP1 is cleaved at the F100∼V101 bond (27), i.e., MMP1 that we used for our experiments has amino acids between V101 and N469. The analysis of the amino acid sequence showed that, between V101 and N469, there are total four CYS residues at C142 (mutated from S142), C278 (native), C366 (mutated from S366), and C466 (native). Note that both active and inactive MMP1 have the same sequence except the E219Q point mutation for inactive MMP1. The analysis of MMP1 crystal structure (4AUO) showed that C278 and C466 are relatively hidden and, therefore, likely to be less labile. Also, we measured the distances between possible combinations of four CYS residues in the open (S142-S366=5.2 nm) and closed (S142-S366=4.5 nm) conformations of MMP1. Only the distance between C142 and C366 changes significantly (∼1 nm) as MMP1 undergoes inter-domain dynamics compared to the changes (∼0.1 nm) in other CYS pair distances. In other words, C278 and C466 would not interfere with smFRET results even if they were labeled. However, there are four possibilities of labeling C142 and C366: 1) no fluorophore attached, 2) only one CYS is labeled, 3) both labeled with the same type of fluorophore, and 4) both labeled with different fluorophores. We would detect smFRET only for the possibility #4 with both C142 and C366 labeled with different dyes. The photophysical properties of fluorophores may also cause problems because fluorescence depends on solution conditions (pH, temperature, chemical composition, etc.) (28). Furthermore, blinking due to transition into metastable states (29, 30) and photobleaching due to reaction with oxygen in the excited state (31) can complicate the tracking of fluorophores. Additionally, the flexible linker between a protein and Alexa dyes (32) and the enzyme microenvironment around the fluorophores can change smFRET (33). Previous studies have elucidated the combined complexity of smFRET, solution conditions, and simulations (34). Since active and inactive MMP1 would be affected equally by these complications, we could distinguish the effects due to enzymatic activity. For further confirmation, we also used inactive MMP1 as the control for all-atom MD simulations (see next paragraph).

### All-atom molecular dynamics simulations

We performed all-atom Molecular Dynamics (MD) simulations using GROMACS version 2018.2 (35). We applied a leapfrog integration time step of 2 fs and constrained all bonded interactions using the LINCS constraint algorithm (36). The short-range neighbor interaction list cut-off was updated every 10 fs and fixed to 0.8 nm. We modeled the short-range interactions using a 12-6 Lennard-Jones potential truncated at 0.8 nm and truncated electrostatic interactions at 0.8 nm. There is no standard choice for truncation. We used 0.8 nm to be consistent with previous publications (37-39). We calculated the long-range electrostatic interactions using the Particle Mesh Ewald scheme (40). We maintained the temperature at 310 K using the Nose-Hoover thermostat (41). We set the pressure at 1 atm using an anisotropic Parrinello-Rahman coupling (42). We defined the simulation cell using periodic boundary conditions in three dimensions and saved the coordinates, velocities, and energies every 5 ps. We used the crystal structure (4AUO) of inactive MMP1 (E219A) bound to model the triple-helical collagen monomer. Note that E219 (27) is the same residue as E200 (12). The number differs because the first residues are different for our full-length MMP1 before activation and the crystal structure 4AUO. The sequence of MMP1 is in the supplementary (**Fig. S1**).

We modified residues in 4AUO to match the amino acid sequences of active and inactive MMP1 used in smFRET experiments using Schrödinger Maestro (Schrödinger Release 2015-4). We incorporated the two serine-to-cysteine (SER142CYS and SER366CYS) substitutions used for labeling MMP1. Additionally, we included an alanine-to-glutamine (A219Q) substitution to match the sequence of inactive MMP1. We repeated procedures in earlier work (37) for 1) initialization of collagen-bound MMP1 complex crystal structure (4AUO), 2) parameterization and implementation steps for the bond distances and angles, and 3) respective force constants and partial charges of the zinc-binding sites.

We placed each protein complex at the center of a cubic unit cell sufficiently large enough to avoid periodic boundary artifacts. We solvated unit cells using TIP4P water molecules and added counterions to neutralize the system. We subjected solvated protein complexes to a steepest descent energy minimization with a maximum force of 1000 kJ/mol stopping criteria for correcting steric clashes between atoms. The minimized structures underwent a 10 ps molecular dynamics simulation using the NVT (fixed particles, volume, and temperature) ensemble with position restraints of 1000 kJ/mol to all heavy atoms. We maintained the position restraints and extended the simulation by 200 ps with constant pressure using the Parrinello-Rahman pressure coupling scheme. We simulated a series of 200 ps intervals, systematically reducing the position restraints to a final 10 kJ/mol. We removed all position restraints with the exception to those on the collagen alpha-carbon atoms, which we increased to 150 kJ/mol to prevent the fraying of the short collagen during the structure relaxation. With restrained collagen, we performed 225 ns NPT (fixed particles, pressure, and temperature) simulation. We removed the position restraints and performed 700 ns NPT simulation without constraints.

### Normal mode analysis using Anisotropic Network Model

We computed preferred conformations or “normal modes” of MMP1 interacting with collagen using the Anisotropic Network Model (ANM) (43). ANM models each amino acid as a bead and creates a virtual bond network connecting these beads (43, 44). By modeling these virtual bonds as harmonic oscillators, ANM calculates normal modes and enables analysis of conformations that are accessible to a 3D protein structure. For the MMP1-collagen system, we used the crystal structure of MMP1 (12) in complex with a collagen peptide (4UAO) using ANM web server 2.1 with default parameters (45).

The output of ANM simulations is an N×N matrix, where N is the number of beads (amino acids) in the system, which provides 3N − 6 non-zero eigenvalues and 3N − 6 eigenvectors that represent the respective frequencies and shapes of the individual normal modes (46). The smaller the frequency, the slower the motion that a mode represents. For each mode, the collective motion of beads and springs is a series of frames. The ANM server allows for the generation of bead coordinates from frames of each mode. We obtained the 3D coordinates of beads from 20 frames of motion for the 20 slowest modes. For each frame, we calculated the desired distances using Pymol (The PyMOL Molecular Graphics System, Version 1.8). We used 1) the distance between S142 and S366 to measure the inter-domain separation, 2) the distance between N171 and T230 to measure the catalytic pocket opening, and 3) the distances between A219 and the cleavage site G∼L (12) to quantify the proximity of MMP1 to the collagen chains.

We obtained the bound configurations of MMP1 domains (for example, see **Fig. 6**) by removing parts from the crystal structure of collagen-bound MMP1 (4AUO). We used Chains A (for MMP1) and C, D, and E (for collagen) as inputs to ANM. We obtained the different lengths of the enzyme (two domains connected by the linker, two domains separated, or the catalytic domain only) by removing the appropriate three-dimensional coordinates from the original PDB file (4AUO): 1) removing no residue, 2) removing the linker region, and 3) removing the hemopexin domain and the linker region. Since 4AUO has amino acids between F100 and C466, we aligned PDB ID 1SU3 (MMP1 structure having amino acids between A32 and C466 with the pro domain) with 4AUO and generated the structure for collagen-bound pro MMP1. ANM is significantly faster than the all-atom MD simulations and provides significant insights. However, there is an intrinsic limitation of the ANM method. While the all-atom MD simulations can distinguish the effect of single point mutations, ANM cannot because each amino acid, regardless of its chemical nature, is modeled as a single bead. As such, a mutation GLU to GLN or GLU to ALA would not have any impact on ANM results if the input structure is the same.

**FIGURE 2.**
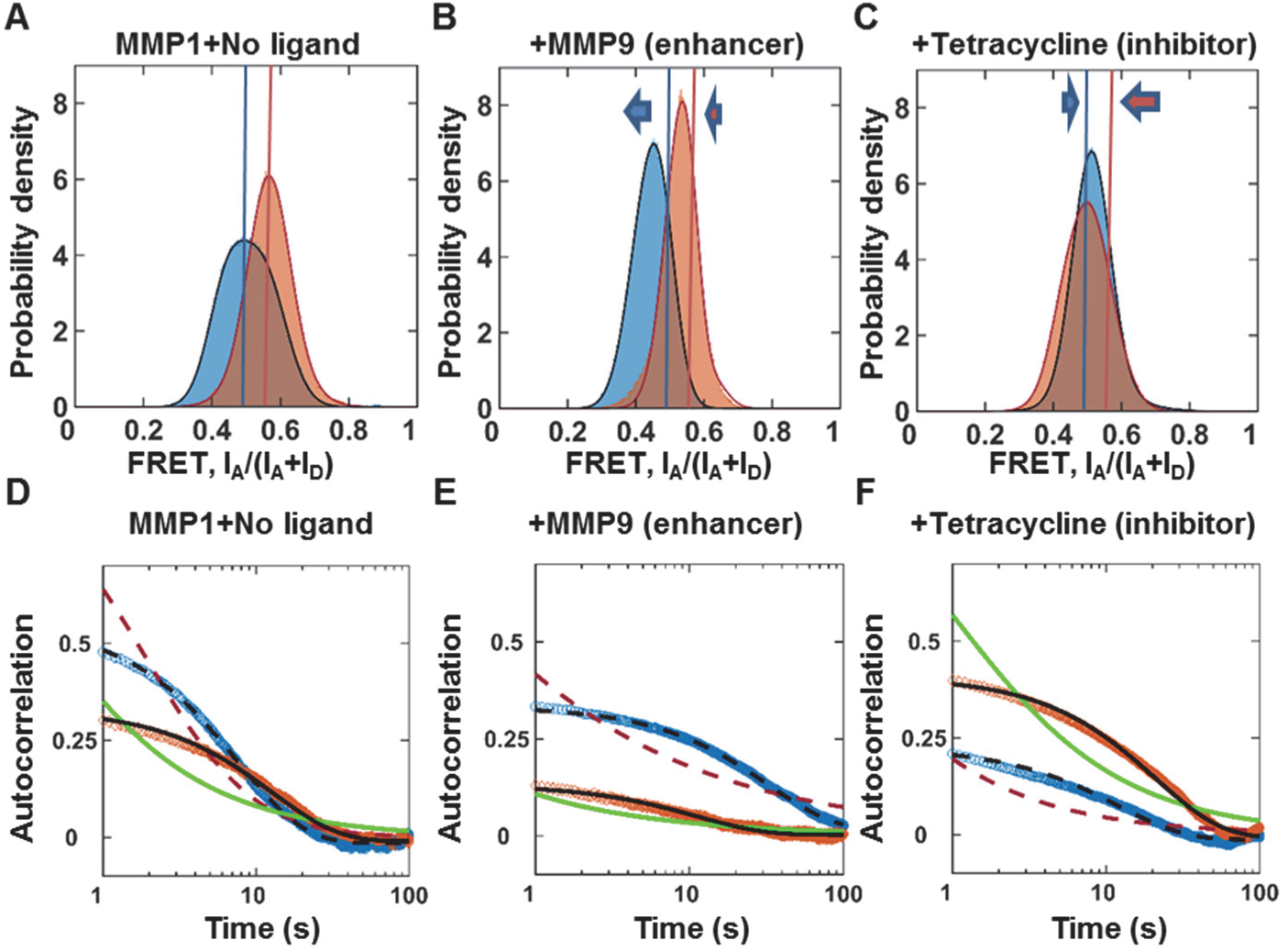
Activity-dependent inter-domain dynamics of MMP1 on reconstituted type-1 collagen fibrils at 22° C with a 100 ms time resolution. Area-normalized histograms of MMP1 inter-domain distance (more than 300,000 FRET values for each condition; bin size=0.005) (**A**) without ligand, (**B**) in the presence of MMP9 (an enhancer), and (**C**) in the presence of tetracycline (an inhibitor) for active (blue) and inactive (orange) MMP1. All histograms are fitted to a sum of two Gaussians (active: solid blue line; inactive: solid red line). Blue and orange lines indicate the peak positions for active and inactive MMP1 without ligands, whereas blue and orange arrows indicate the directions of shifts of the FRET peaks in the presence of ligands. Autocorrelations of MMP1 inter-domain distance (**D**) without ligand, (**E**) in the presence of MMP9, and (**F**) in the presence of tetracycline for active (blue) and inactive (orange) MMP1. All autocorrelations are fitted to exponentials and power laws (exponential fit to active: dashed black line; power law fit to active: dashed red line; exponential fit to inactive: solid black line; power law fit to inactive: solid green line). The error bars in histograms and autocorrelations represent the square roots of the bin counts and the standard errors of mean (sem) and are too small to be seen. The supplementary information contains the fit equations and the best-fit parameters for histograms (**Table S1**) and autocorrelations (**Table S2**).

**FIGURE 3.**
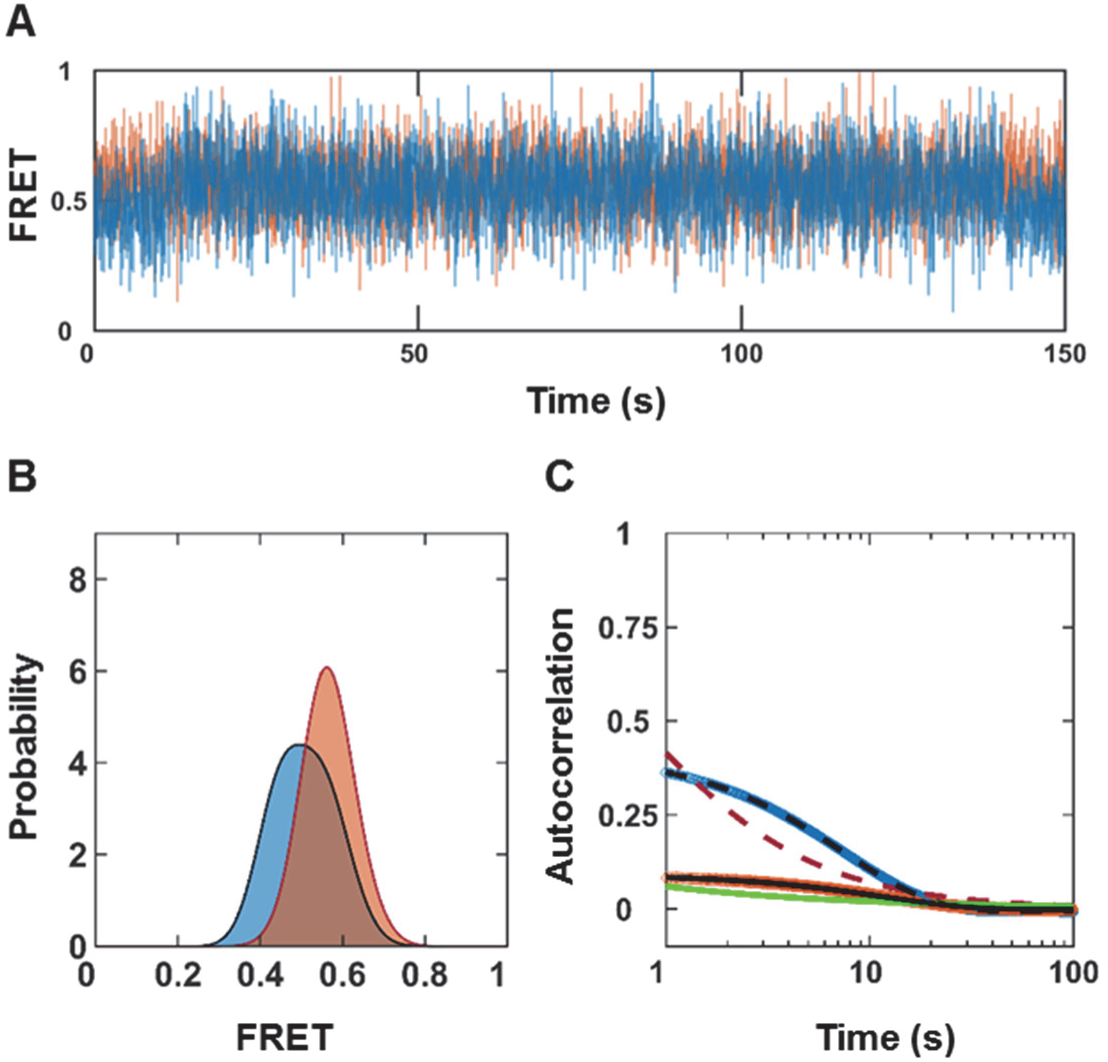
Simulated MMP1 inter-domain dynamics as a Poisson process. (**A**) Examples of simulated two-state FRET trajectories with noise for active (blue) and inactive (orange) MMP1. (**B**) Histograms of simulated FRET values. (**C**) Autocorrelations of simulated trajectories recover the sum, k1+k2, from exponential fits (active: dashed black line; inactive: solid black line). As expected, power law does not fit autocorrelations (active: dashed red line; inactive: solid green line). Exponential fits recover k1+k2 with and without noise. The addition of noise changes the width of the FRET histograms and y-intercepts. The error bars are the sems for histograms and autocorrelations and are too small to be seen. The supplementary information contains the best-fit parameters for histograms (**Table S4**) and autocorrelations (**Table S5**).

**FIGURE 4.**
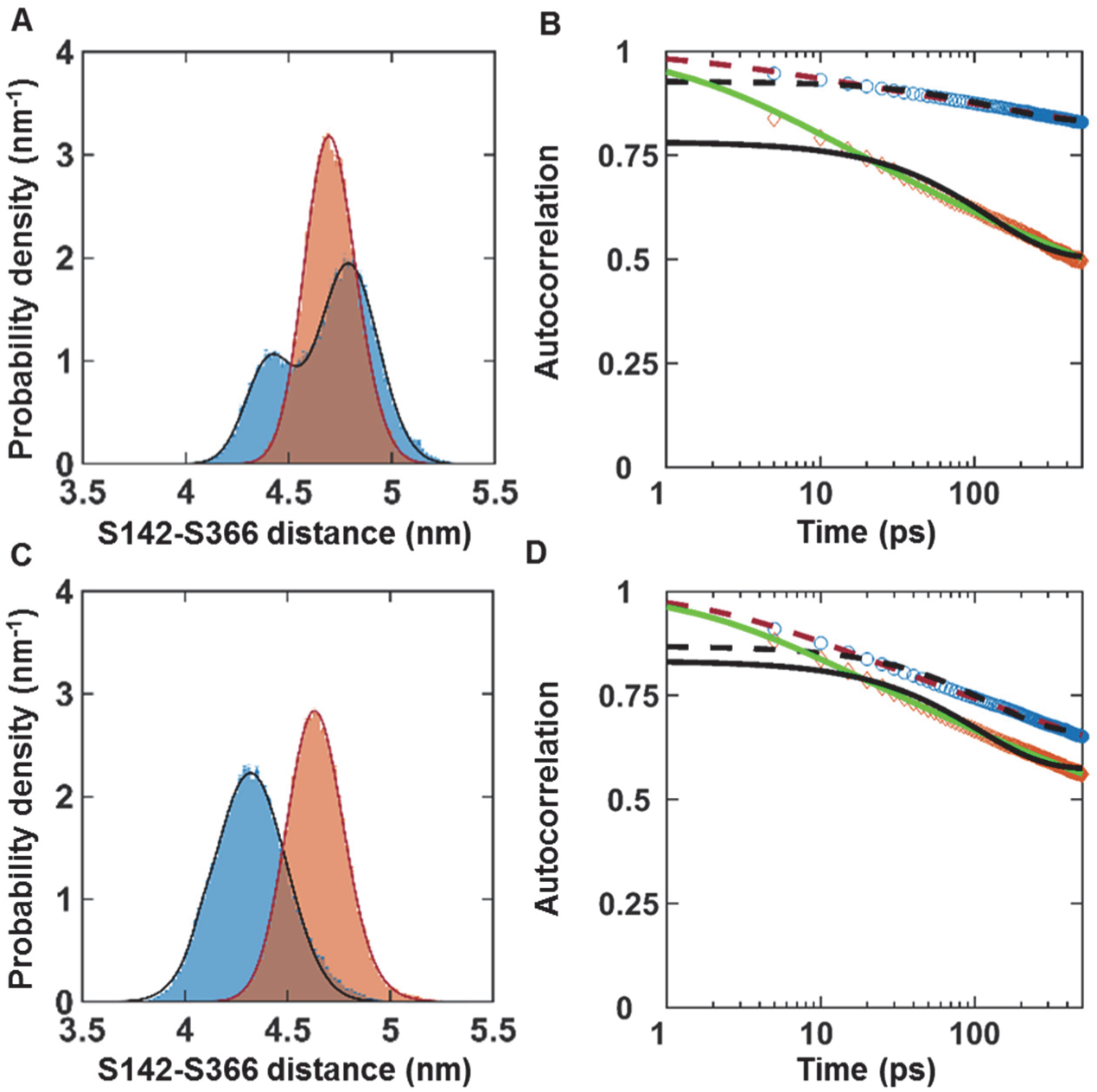
All-atom MD simulations with collagen backbone restrained (A and B) and unrestrained (C and D). Simulated dynamics using GROMACS simulation package at 37° C with 2 fs time step; data saved every 5 ps; 225 ns and 700 ns simulations for restrained and unrestrained, respectively. (**A**) Area-normalized histograms of simulated dynamics with bin size=0.02 nm and (**B**) autocorrelations for active (blue) and inactive (orange) with the collagen backbone restrained by an energy penalty of 1000 kJ/mol. (**C**) Area-normalized histograms of simulated dynamics and (**D**) autocorrelations for active (blue) and inactive (orange) with the collagen backbone unrestrained. All histograms are fitted to a sum of two Gaussians (active: solid blue line; inactive: solid red line). All autocorrelations are fitted to exponentials and power laws (exponential fit to active: dashed black line; power law fit to active: dashed red line; exponential fit to inactive: solid black line; power law fit to inactive: solid green line). The error bars in the histograms represent the square roots of the bin counts and are too small to be seen, whereas the autocorrelations do not have error bars. The supplementary information contains the best-fit parameters for histograms (**Table S6**) and autocorrelations (**Table S7**).

**FIGURE 5.**
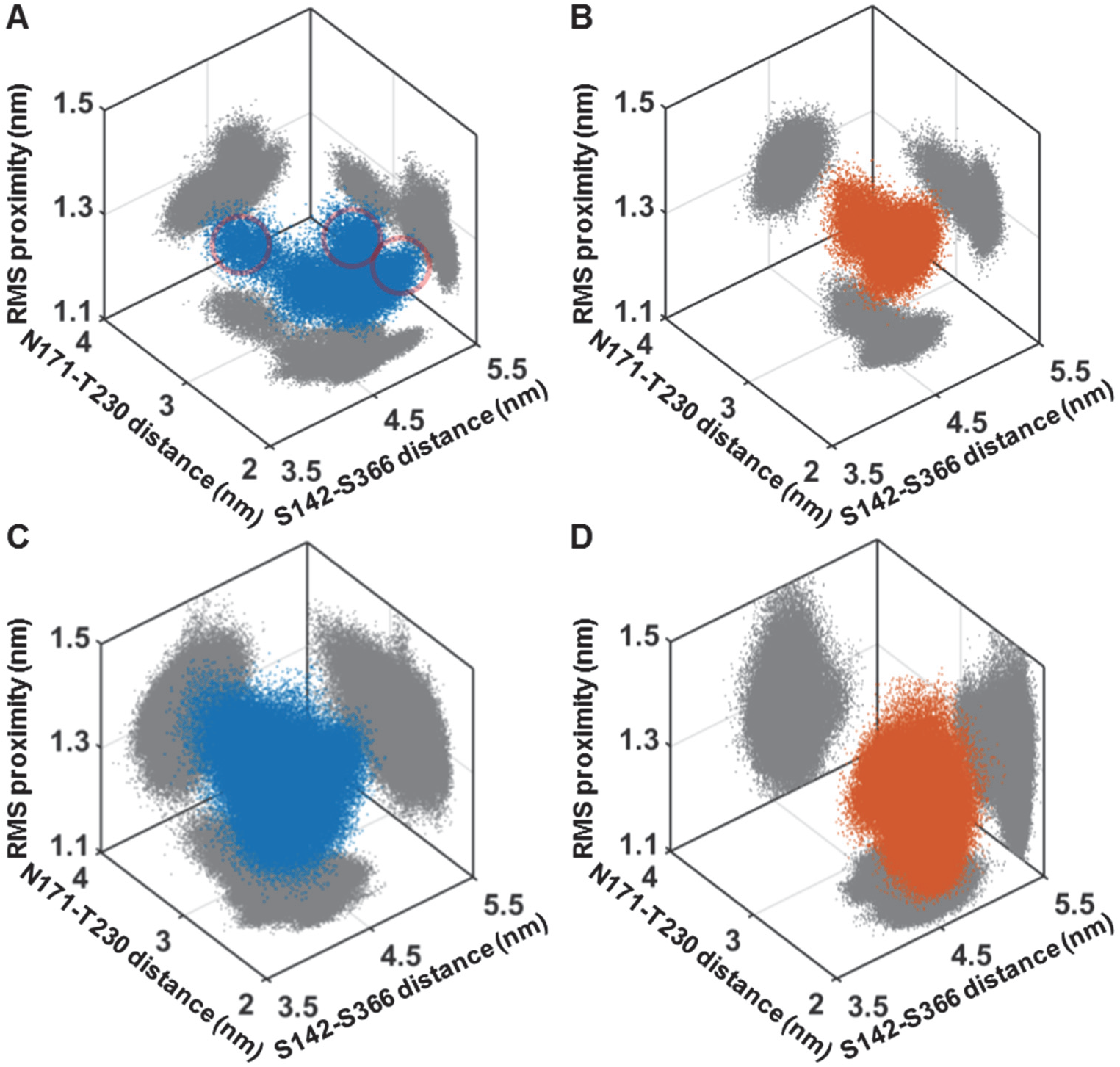
Insights from all-atom MD simulations with collagen backbone restrained (A and B) and unrestrained (C and D). Three-dimensional scatter plots of S142-S366 distance (represents inter-domain dynamics), N171-T230 distance (represents the opening of the MMP1 catalytic pocket), and rms distance between the MMP1 catalytic site and the cleavage sites on three collagen chains for active (blue) and inactive (orange) MMP1. Two-dimensional projections of the scatter plots are in gray. The clusters encircled in red in (**A**) represent the plausible catalytically relevant conformations.

**FIGURE 6.**
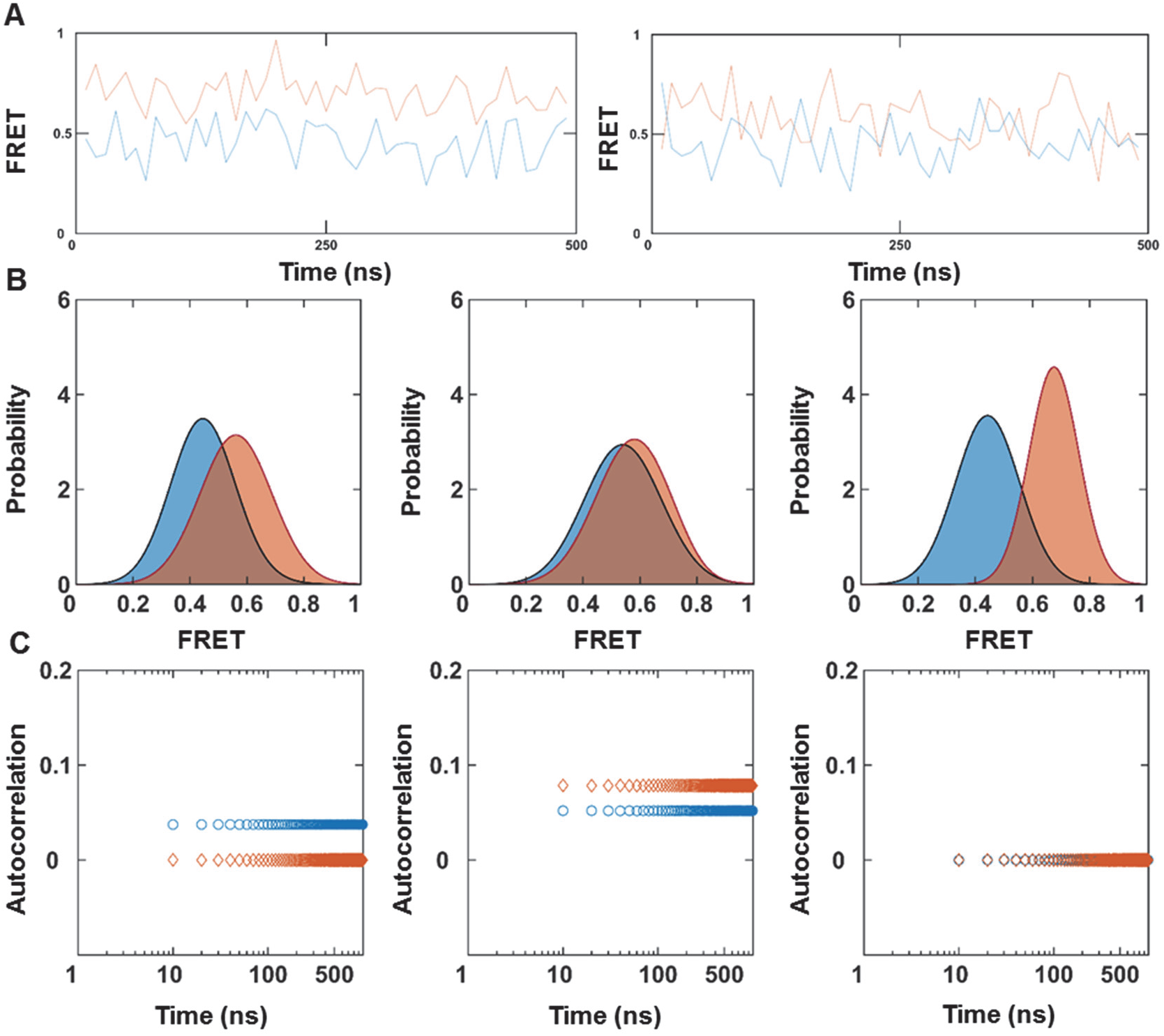
Two-state simulations at faster nanosecond timescales. Different trials with the same underlying parameters produced different (**A**) trajectories (blue: active MMP1; orange: inactive MMP1), (**B**) histograms with bin size=0.005 (active: solid blue line; inactive: solid orange line), and (**C**) autocorrelations (active: blue circle; inactive: orange diamond). Different trials of simulations lead to different results even though the underlying process is the same two-state Poisson process. It is challenging to see hidden transitions in the example trajectories because the difference between S1 and S2 (∼0.1) is similar to the noise levels (∼0.1). For simulated trajectories with the same parameters but without noise, see **Fig. S3 A**.

## Results and discussion

### Single-molecule measurement of MMP1 inter-domain dynamics on reconstituted type-1 collagen fibril

For our studies, we differentiated the role of activity by comparing the results for active (E219) with catalytically inactive (E219Q) MMP1. We mutated SER142 in the catalytic domain and SER366 in the hemopexin domain to cysteines (**Fig. 1 A**). Note that the residue E219 (27) is the same as E200 (12), which differs due to the first residue used for counting. The distance between the central carbon atoms of the two selected amino acids is ∼4.5 nm, which is similar to the Förster radius of 5.1 nm (47) for the dye pair Alexa Fluor® 555 and Alexa Fluor® 647. We labeled MMP1 with Alexa555 and Alexa647 using maleimide chemistry. The left panel of **Fig. 1 B** shows the gel electrophoresis of labeled MMP1. The right panel of **Fig. 1 B** presents the specific activities of labeled and unlabeled MMP1 on the synthetic peptide substrate, MCA-Lys-Pro-Leu-Gly-Leu-DPA-Ala-Arg-NH_2_ (48). Labeling did not affect the function of MMP1. We used a TIRF microscope because many MMPs can be imaged simultaneously, in contrast to a confocal microscope with point detection. We used a TIRF microscope in a prism-type configuration (**Fig. 1 C**) because it has lower background noise due to the separation of excitation and emission paths, in contrast to the objective-type TIRF microscope where the excitation and emission paths share optics. As labeled MMPs moved and underwent inter-domain dynamics on the surface of a fibril, fluorescence emissions from the donor and acceptor dyes were detected using two quadrants (channels) of an EMCCD camera. We superimposed the two channels and tracked the intensity and location of spots to measure the inter-domain dynamics. When the two domains are within the Forster radius, the energy from Alexa555 is non-radiatively transferred to Alexa647 due to FRET. As a result, the fluorescence from Alexa555 decreases leading to a simultaneous increase in the emission from Alexa647. **Fig. 1 D** shows an example of anticorrelated fluorescence from the two dyes due to MMP1 inter-domain dynamics. See methods for detailed procedures.

### Well-separated catalytic and hemopexin domains correlate with the activity of MMP1

The role of dynamics in function is not clear: Warshel proposed that protein dynamics and enzyme catalysis are not coupled (2, 49), but Karplus proposed that enzyme motions can lower the activation energy (1, 3). We measured the inter-domain dynamics of active and inactive MMP1 on reconstituted type-1 collagen fibrils using smFRET (**Fig. 2**). The histograms of FRET values revealed the equilibrium distributions of inter-domain distance, whereas the autocorrelations showed how conformations correlate with each other at different time points. We fitted a sum of two Gaussians to the histograms (**Fig. 2 A**). The best-fit parameters for the centers of Gaussians are: 0.44±0.01 and 0.55±0.01 for active MMP1; 0.56±0.01 and 0.68±0.01 for inactive MMP1 (**Table S1**). These results suggest that the low FRET conformation at 0.44±0.01, where the two MMP1 domains are distant (the open conformation), are relevant for the catalytic activity of MMP1. The relevance of the open structure in function is in agreement with previous results for collagen monomers based on NMR and SAXS experiments and all-atom MD simulations (13, 15-17) that revealed the importance of a larger separation between the catalytic and hemopexin domains for activity. As such, the open conformation with more inter-domain distance is functionally relevant for MMP1 interactions with both collagen monomers and fibrils.

To further delineate the relation between inter-domain dynamics and activity, we used two ligands that modulate MMP1 activity: MMP9 (an enhancer) and tetracycline (an inhibitor). MMP9, another member of the MMP family, is generally thought to be unable to degrade structurally intact triple-helical type-1 collagen (50). Both pro (51, 52) and activated forms (53-56) of MMP9 exist in numerous physiological conditions. MMP9 and MMP1 form a stable complex (24), and computational studies have predicted enhancement of MMP1 activity (57). Tetracycline plays dual roles as an antibiotic and an MMP inhibitor (23). Our measurements confirmed that MMP9 is an enhancer, whereas tetracycline is an inhibitor of MMP1 activity (**Fig. S2 I**). The surface morphology of fibrils treated with MMP1 changed in the presence of MMP9 and tetracycline (**Fig. S2 A-H**).

In the presence of MMP9, the peaks shift towards the left to 0.43±0.01 and 0.48±0.01 for active MMP1 (**Fig. 2 B**). In contrast, tetracycline inhibits low FRET conformations and appears to move the peaks towards the right to 0.51±0.01 and 0.55±0.01 for active MMP1 (**Fig. 2 C**). In other words, MMP1 stays more or less in the open conformation (low FRET) as its activity increases or decreases in the presence of MMP9 and tetracycline, respectively. In contrast, inactive MMP1 primarily stays in a closed conformation corresponding to FRET values at ∼0.50 or higher (**Table S1**). In the presence of tetracycline, both active and inactive MMP1 primarily adopt conformations with FRET values at ∼0.50 (**Table S1**). We performed molecular docking to investigate why tetracycline affected active and inactive MMP1 similarly. We observed that tetracycline forms two hydrogen bonds with each domain (**Fig. S4**) and appears to hold MMP1 in place, preventing inter-domain dynamics. As such, the molecular docking provides an explanation for similar FRET values for active and inactive MMP1 in the presence of tetracycline. These results on activity-dependent inter-domain dynamics collectively confirm that open MMP1 conformation and catalytic activity of MMP1 are functionally related.

Next, we studied correlation in the dynamics because correlated motions indicate a decrease in conformational entropy. A reduction in entropy may affect kinetics and thermodynamics of biological processes, including catalysis (58). Without any ligand, active MMP1 had more time-correlated conformational dynamics at short times than inactive MMP1 (**Fig. 2 D**). In the presence of MMP9, the time-correlations were longer for both active and inactive MMP1 (**Fig. 2 E**). MMP9 not only led to more low FRET conformations of MMP1, but it also stabilized low FRET conformations as indicated by narrow widths of histograms (**Fig. 2 B** and **Table S1**) and longer correlation times (**Fig. 2 E** and **Table S2**). Interestingly, tetracycline reversed the trend, and inactive MMP1 had longer time-correlation than active MMP1 (**Fig. 2 F** and **Table S2**). In other words, ligands can alter conformational entropy of MMP1, leading to a change in function.

The observed modulation of inter-domain separation and correlation could be a “consequence” or the “cause” of catalytic activity. We argue that low FRET conformations are likely the “cause” for several reasons. First, a mutation E219Q at the catalytic site not only renders MMP1 inactive, but it also induces changes in the distance allosterically to alter the inter-domain dynamics (**Fig. S5**). Second, changes in the linker region also have been shown to affect activity allosterically (59). Third, the catalytic domains of MMPs are mostly similar (7), suggesting a crucial role of the hemopexin domains in substrate specificity (60). Fourth, the catalytic domain alone can degrade denatured collagen, where the cleavage sites are easily accessible. However, the catalytic domain requires the hemopexin domain to degrade the basic building block of collagen fibrils, i.e., triple-helical collagen, where the cleavage sites are hidden (4, 61, 62). Overall, it appears that the hemopexin domain communicates with the catalytic domain and induces necessary changes to enable collagen degradation. Further studies are required to establish causality.

### MMP1 inter-domain dynamics is a two-state Poisson process

MMP1 inter-domain dynamics are random (stochastic) time series, which can be described by a Poisson process or a set of Poisson processes. A Poisson process has a constant probability of occurring at each temporal or spatial point. Previously, we applied the Poisson process approach to diverse phenomena (63), including MMP1-collagen interactions (18, 19). To investigate if the Poisson process approach could be applied to describe MMP1 inter-domain dynamics, we performed stochastic time-series simulations of the distance between SER142CYS and SER366CYS. We analyzed simulations and experimental FRET measurements similarly.

The centers of the Gaussians fits (**Table S1**, the parameters b1 and b2) were defined to be the two states, S1 (lower FRET) and S2 (higher FRET). We considered that MMP1 interconverts between S1 and S2 with kinetic rates k1 (S1⟶S2) and k2 (S2⟶S1). We calculated the ratio k1/k2 from the ratio Area(S2)/Area(S1) determined by the Gaussian fits to FRET histograms (**Fig. 2 A-C**). We derived the sum k1+k2 from the decay rate e (**Table S2**) determined by the exponential fits (**Fig. 2 D-E**) because the autocorrelation for a two-state system decays as the sum of two rates (64). The ratio k1/k2 and sum k1+k2 determined k1 and k2 (**Table S3**) for different experimental conditions. We considered that if MMP1 is in state S1, there is a constant probability, given by P(t)dt=k1*exp(-k1*dt)*dt, of going to state S2 at each time step dt. Similarly, if MMP1 is in state S2, there is a constant probability, given by P(t)dt=k2*exp(-k2*dt)*dt, of going to state S1 at each time step dt. To implement this kinetic mechanism, we used the “exprnd” function in Matlab to generate the exponentially distributed dwell-times in each state with and without noise.

As an example, we considered the inter-domain dynamics of MMP1 without ligands and simulated 350 FRET trajectories with (**Fig. 3**) and without (**Fig. S3**) noise with 100 ms time resolution, each 1000 s long. We added Gaussian-distributed noise to the simulated FRET trajectories determined by the Gaussian fit parameters (**Table S1**) for the experimental histograms (**Fig. 2 A-C**). **Fig. S3** shows examples of FRET trajectories for active and inactive MMP1 without noise (**Fig. S3 A**), normalized histograms (**Fig. S3 B**), and autocorrelations (**Fig. S3 C**). The histograms of simulated FRET values reproduced the locations of states (**Table S4**), and an exponential fit the autocorrelations than power laws as expected. The analysis of simulated trajectories recovered the experimentally-determined input rates for both active and inactive MMP1 without noise (**Table S5**). We also simulated FRET trajectories using the same parameters but with noise. **Fig. 3** shows examples of FRET trajectories with noise for active and inactive MMP1 (**Fig. S3 A**), normalized histograms (**Fig. 3 B**), and autocorrelations (**Fig. 3 C**). Even with noise, Gaussian fits recovered the simulated FRET values (**Table S4**) and the ratio of rates, whereas exponential fits recovered the sum of decay rates (**Table S5**). Note that the y-intercept of autocorrelation depends on S1, S2, k1, k2, and noise. Without noise, a closed-form expression for the y-intercept can be derived (64). In summary, MMP1’s inter-domain dynamics can be described by a Poisson process at the experimental timescales.

A two-state Poisson process description of MMP1 inter-domain dynamics enables a more in-depth interpretation of autocorrelations (**Fig. 2 D-E**) and exponential fit parameters (**Table S3**). For example, if low FRET values at ∼0.44±0.01 for active MMP1 with and without MMP9 represent catalytically relevant open conformations, we would expect MMP1 to stay in this state longer as the activity increases. In other words, MMP9 would facilitate MMP1 staying in the low FRET state longer, leading to a smaller k1, which is indeed the case as indicated by the decrease of k1 from 0.08 to 0.01 s^−1^ in the presence of MMP9 (**Table S3**). In other words, the kinetic rates of interconversion between open and closed MMP1 conformations correlate with the activity.

### A larger inter-domain distance often accompanies a larger catalytic pocket opening of MMP1 and closer proximity to the collagen chains

The MMP1 catalytic cleft (∼0.5 nm wide) is too narrow to accommodate the collagen monomer (1.5 nm in diameter) (13). Therefore, a larger opening of the MMP1 catalytic pocket is needed to bring collagen closer to the active site and must accompany more inter-domain separation if low FRET conformations are indeed relevant. To this end, we investigated the activity-dependent inter-domain dynamics of active and inactive MMP1 using all-atom simulations (see methods).

As shown in **Fig. 4 A**, the all-atom simulations reproduced that active MMP1 has more low FRET conformations (i.e., inter-domain distances represented by the blue shoulder around 5 nm) with the collagen backbone restrained. In contrast, MMP1 has fewer low FRET states with the collagen backbone unrestrained (**Fig. 4 C**). **Table S6** contains the best-fit parameters for Gaussian fits to the histograms. The autocorrelations (**Fig. 4 B** and **D**) of simulated inter-domain distances show that active MMP1 has more time-correlated dynamics at short times than inactive MMP1 in agreement with the experiments. Note that an exponential fit autocorrelations better than power law for smFRET measurements (**Fig. 2 D-F**). In contrast, power law fits autocorrelations better than an exponential for simulations (**Fig. 4 B** and **D**). The best-fit parameters for the power law (*a* × *t* +1)^−*b*^ are in **Table S7**. The error bars are the sems. There are reports on power law distribution of conformational dynamics (65, 66). However, we need further studies to understand the transition from power law to an exponential distribution at the two extreme timescales.

For unrestrained collagen backbone, the histograms of simulated conformations (**Fig. 4 C**) showed that active MMP1 adopts more high FRET conformations (smaller inter-domain distances), in contrast to the experimental observations that active MMP1 adopts more low FRET conformations (**Fig. 2 A**). In other words, the all-atom simulations with a restrained collagen backbone agree more with the experiments, which may be explained by the fact that the organization of fibrils restrains collagen monomers in comparison to collagen monomers in solution. Also, the difference in autocorrelations for active and inactive MMP1 on unrestrained collagen (**Fig. 4 D**) is significantly lower than those on restrained collagen (**Fig. 4 B**). Furthermore, simulations also suggest that MMP1 dynamics vary depending on the substrate properties, i.e., enzyme-substrate interactions are mutually affected by each other. Thus, experiments and simulations provide a synergistic combination to gain insights into MMP1 activity. Informed by single-molecule measurements of MMP1 inter-domain dynamics, we used MD simulations to identify catalytically relevant conformations more precisely. We assumed that the catalytic pocket should open more before catalysis so that the catalytic residues can approach the cleavage sites on the three collagen chains. We measured the catalytic pocket opening by the distance between N171 and T230. We quantified the proximity to the collagen chains by the rms distance between the MMP1 residue (E/Q219) and the collagen cleavage sites (G∼L).

Three-dimensional scatter plots (**Fig. 5**) show the patterns in the MMP1 conformational landscape. **Fig. 5 A** suggests that the encircled clusters are likely the functionally relevant conformations of MMP1 because these conformations are significantly absent in inactive MMP1 (**Fig. 5 B**). When the collagen backbone is free, the detailed structures of the conformational space disappear (**Fig. 5 C-D**). Interestingly, the number of conformational peaks depends on the projection plane (reaction coordinates). Overall, we compared simulations with experiments to gain further insights into the MMP1-collagen system and learned that an increase in inter-domain separation correlates with a larger catalytic pocket opening and closer proximity to the collagen chains.

The all-atom MD simulations also explained why the E219Q mutation in the catalytic domain alters inter-domain dynamics in the hemopexin domain. We calculated the root-mean-squared side-chain fluctuation (RMSF) for residues in active and inactive MMP1 (**Fig. S5**). The E219Q mutation caused significant changes in fluctuations allosterically in the hemopexin domain. Surprisingly, variations in the catalytic domain are similar for active and inactive MMP1. The means±stds of RMSF (**Fig. S5**) across all the amino acids are 3.6±0.7 nm for active and 4.6±0.4 nm for catalytically inactive MMP1. The higher std of RMSF for active MMP1 might explain the broader widths of histograms for active MMP1. To further delineate the difference between active and inactive MMP1, we performed principal component analysis (PCA) of the fluctuations (67) (see supplementary information). We observed a significant difference between active and inactive MMP1 (**Fig. S6, Fig. S7**, and **Fig. S8**).

### MMP1 undergoes substrate-dependent adaptive conformational dynamics

Both the experiments (**Fig. 2**) on collagen fibrils at 100 ms timescale and the simulations (**Fig. 4**) with restrained collagen monomer at 5 ps timescale showed that active MMP1 adopts more conformations with larger inter-domain distances. Such changes in protein conformations can occur due to interactions with a ligand/substrate and can be understood based on the general principles of classical or quantum mechanics. If the interactions are weak, we can still use the conformations proteins without interactions and treat any conformational changes as perturbations. For weak interactions, one can envision applying “lock-and-key” (68), “induced fit” (69), and “conformational selection” (70) theories of enzyme specificity and function (71). If the interactions are strong, however, conformations/normal modes/eigenfunctions are not represented by those of individual systems. For the MMP1-collagen system, the interactions are so strong that there is negligible unbinding once MMP1 binds collagen (19, 21). As a result, collagen-binding can lead to a new set of allowed MMP1 conformations. Indeed, MMP1 undergoes conformational dynamics in a substrate-dependent manner because restraining the collagen backbone changes the structures and correlations (**Fig. 4**). Besides, the mutation E219Q at the catalytic site of MMP1 also changes the set of allowed conformations that appear at the two extreme timescales of the all-atom simulations (**Fig. 3** and **Fig. 4**) and experiments (**Fig. 2**). Therefore, we can propose an alternative model of “adaptive conformational dynamics” where substrate recognition leads to a new set of protein conformations that can be dynamically influenced by mutations and ligands (**Fig. 2**). It is also plausible to consider adaptive conformational dynamics as time-dependent disorder or randomness, which is known to alter chemical rates (72).

### Poisson process connects experimental and all-atom simulation timescales

The agreements between experiments and simulations suggest similar initial conformations of collagen-bound MMP1 at picoseconds as well as milliseconds. Note that crystal structures represent thermodynamically (lowest energy) or kinetically (lower activation barriers) states (73). Since we used crystal structures for simulations and we detected collagen-bound MMP1 for smFRET measurements, similar initial conformations are plausible. As such, we postulated that functionally relevant MMP1 inter-domain conformations have a finite probability of occurring even at picoseconds, and dynamics might be connected using a Poisson process approach.

We modeled MMP1 inter-domain dynamics as a two-state Poisson process, and Poisson process-based stochastic simulations reproduced the kinetics of interconversion. In principle, a Poisson process at second timescales of experiments should also be a Poisson process at picosecond timescales of simulations. However, the probability of transition between states will be significantly lower at picosecond according to the equation for probability for the exponential distribution P(t)dt=k*exp(-k*dt)*dt. For example, k is around 0.1 s^−1^ for MMP1 (**Table S3**), and chances for a transition at each time step are 0.0099 and 5*10^−13^ for 100 ms (smFRET experiments) and 5 ps (all-atom MD simulations) time steps, respectively. As a result, stochastic simulations at second timescales are more likely to have transitions and would reproduce the experimental features. However, simulations at ps-ns timescales are less probable to have interconversion during the length of simulated trajectories and would give different results at different trials (**Fig. 6**) and may not reproduce the correlations even if the underlying process is a Poisson process.

To determine if we could apply the Poisson process approach to the faster timescales of MD simulations, we simulated 1000 ns long trajectories with time resolution, dt=10 ns, using the same experimentally determined parameters, S1, S2, k1, and k2 for MMP1 without ligands (**Table S1** and **Table S3**). As shown in **Fig. 6**, both histograms and autocorrelations show different results at different trials because of the statistical nature of transitions. Without transitions, a trajectory is simply a constant value (S1 or S2) plus noise leading to zero autocorrelations (**Fig. 6**). If there are transitions, autocorrelations appear (**Fig. 6**).

The power law behavior of correlations (74) at picosecond timescales is intriguing because it usually occurs when there is long-range order (for example, phase transitions (74, 75)) or correlated random walk (for example, stock market fluctuations (76) and DNA sequences (77, 78)). We have several possible explanations. First, an exponential can become linear for short timescales if the condition kt ≪ 1 is valid. Second, two-state Poisson process simulations showed that exponential correlations (**Fig. 3, C**) in the presence of noise changed into power law correlations in the absence of noise (**Fig. S3, C**) for the same underlying parameters. Indeed, fluctuations can transform a Lorentzian line shape into a Gaussian line shape (79). Variations in measurements can originate from samples, instruments, and tracking methods, leading to changes in the observed pattern (80). Third, **Fig. 6** also provides one explanation for the power law behavior at short timescales because the presence of correlations stochastically depends on the number of transitions during the simulated length of trajectories. Without any transition, the correlation is zero. With transitions, the correlation is not zero but does not decay in ps-ns timescales because the rates of conversion between the two states (∼0.1 s^−1^) are slow. As a result, the simulated correlation fits better with a power law. Further studies are needed to explain the power law behavior of autocorrelations at picosecond timescales.

### Both the linker and the collagen substrate facilitate inter-domain communications

We computed preferred conformations or “normal modes” of collagen-bound MMP1 using the ANM simulation (43) (**Fig. 7**). The ANM is a prediction method that models each amino acid as a bead and creates a virtual bond network that connects these beads based on a user-defined cut-off distance (43, 44). By modeling these virtual bonds as harmonic oscillators, we obtained an integrated view of the restrictions in conformational space experienced by each bead and, in turn, predicted what motions are accessible to MMP1. ANM cannot distinguish between active and inactive MMP1. Since active MMP1 with the pro domain attached (proMMP1) is also catalytically inactive, we calculated normal modes for proMMP1 as a proxy for inactive MMP1. Note that we produced both active (E219) and inactive (E219Q mutant) MMP1 as pro enzymes with the pro domain attached (27). MMPs are catalytically inactive with the pro domains attached. The pro domains need to be cleaved to activate MMPs. We activated proMMP1 using trypsin, as described in our previous publication (27). For ANM simulations, we used activated (pro domain cleaved) and pro (pro domain attached) MMP1. With the pro domain attached, we expected that the catalytic pocket opening and proximity to the collagen chains for proMMP1 would be restricted.

**FIGURE 7.**
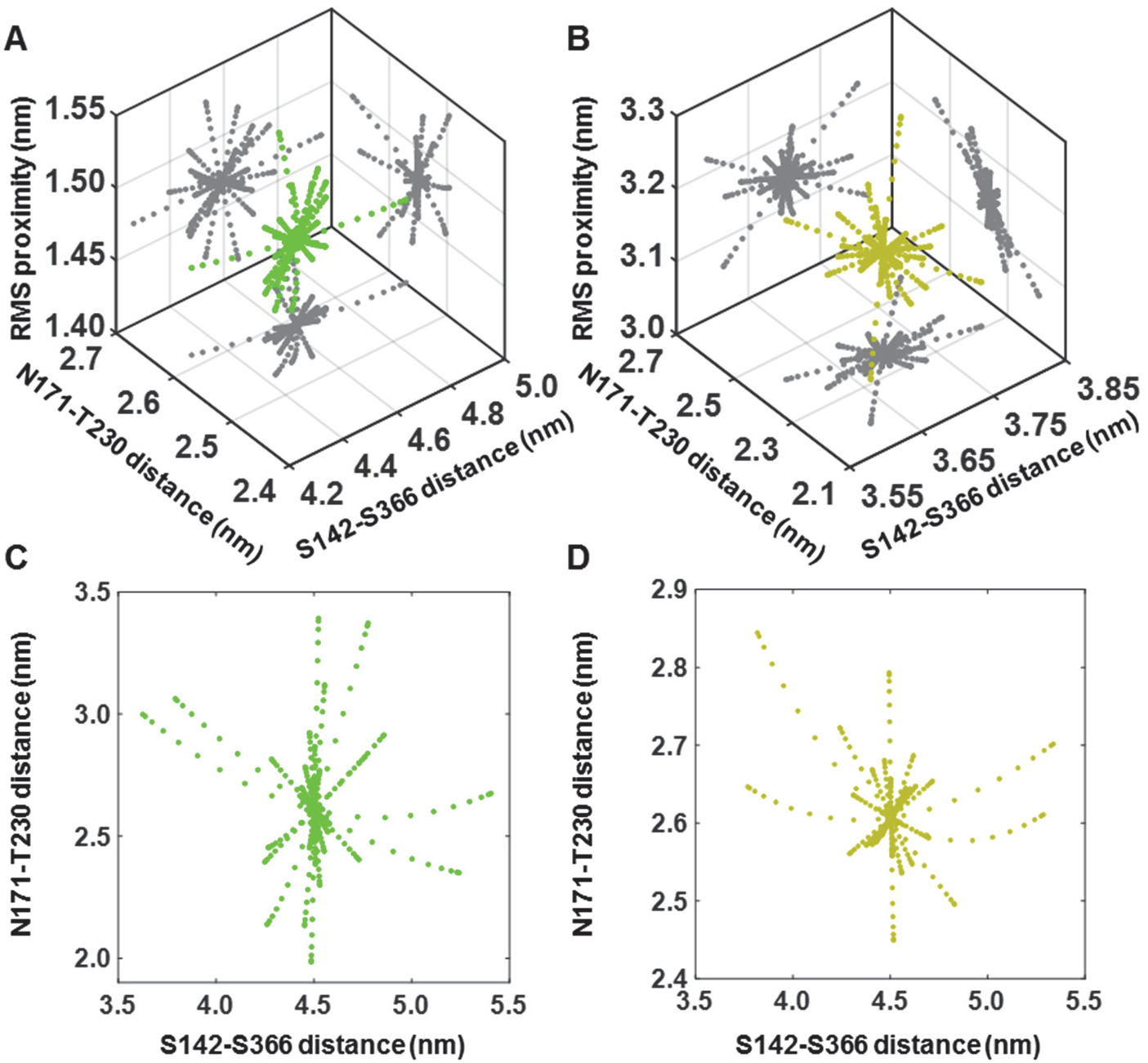
Normal mode analysis of MMP1 using the Anisotropic Network Model. Three-dimensional scatter plots of S142-S366 distance (represents inter-domain dynamics), N171-T230 distance (represents the opening of the MMP1 catalytic pocket), and rms distance between the MMP1 catalytic site and the cleavage sites on three collagen chains for (**A**) collagen-bound activated (green) and (**B**) pro (greenish-yellow) MMP1. Two-dimensional projections in **A** and **B** of the scatter plots are in gray. Two-dimensional scatter plots of S142-S366 distance and N171-T230 distance for free (**C**) activated (green) and (**D**) pro (greenish-yellow) MMP1.

We calculated normal modes for collagen-bound (**Fig. 7 A** and **Fig. 7 B**) and free (**Fig. 7 C** and **Fig. 7 D**) MMP1 for activated and pro MMP1. As shown in **Fig. 7**, free and collagen-bound MMP1 have some similarities in allowed normal modes. Some modes of free MMP1 disappear, and some new modes appear for collagen-bound MMP1. When MMP1 is bound to collagen, the inter-domain distance between S142 and S366 has larger values for activated MMP1 than pro MMP1. The larger inter-domain distances accompanied larger catalytic pocket openings between N171 and T230 and a smaller rms proximity to the collagen cleavage sites for activated MMP1 (**Fig. 7 A**) in comparison to pro MMP1 (**Fig. 7 B**). When MMP1 was not bound to collagen, the ANM calculations showed that the catalytic pocket openings have a broader range for activated MMP1 (**Fig. 7 C**) in comparison to pro MMP1 (**Fig. 7 D**), but the inter-domain distances are similar. These results further confirm that the collagen substrate can significantly change the dynamics of MMP1.

We investigated whether both the linker and collagen can allosterically communicate between the two domains. **Fig. 8** shows that the catalytic domain opening has the largest std (0.52 nm) when both the linker and collagen are present (**Fig. 8 A**). The catalytic pocket opening has the smallest std (0.19 nm) when only the catalytic domain interacts with collagen (**Fig. 8 C**). The changes in opening suggest the essential role of the hemopexin domain in MMP1 activity. We computed conformations without the linker and observed that the hemopexin domain can still influence the catalytic domain plausibly via collagen and increase the catalytic pocket opening (**Fig. 8 B**). In this case, the catalytic pocket opening has an std of 0.26 nm, which is between 0.52 nm and 0.19 nm (**Fig. 8**). In other words, the two MMP1 domains can communicate via collagen even when the linker does not physically connect the two domains. Previous studies reported that a mixture of the two MMP1 domains purified separately could degrade triple-helical collagen (4), but the reason was not apparent. A larger catalytic opening due to allosteric communications between the two MMP1 domains via collagen explains why even a mixture of the two MMP1 domains can degrade triple-helical collagen. Such allosteric communications may lead to the observed correlations in inter-domain motions (**Fig. 2 D-F**). Note that both autocorrelation and entropy are measures of randomness and order (81). A higher correlation suggests a lower amount of randomness and a higher amount of order. The randomness and order also change entropy that connects to kinetics and thermodynamics. In other words, any correlated motion or stabilization of conformations due to allosteric communications indicates a decrease in randomness (entropy). Enabling access to the cleavage sites on triple-helical collagen via inter-domain dynamics of MMP1 is an entropy-driven process, which can lead to an increased rate of cleavage.

**FIGURE 8.**
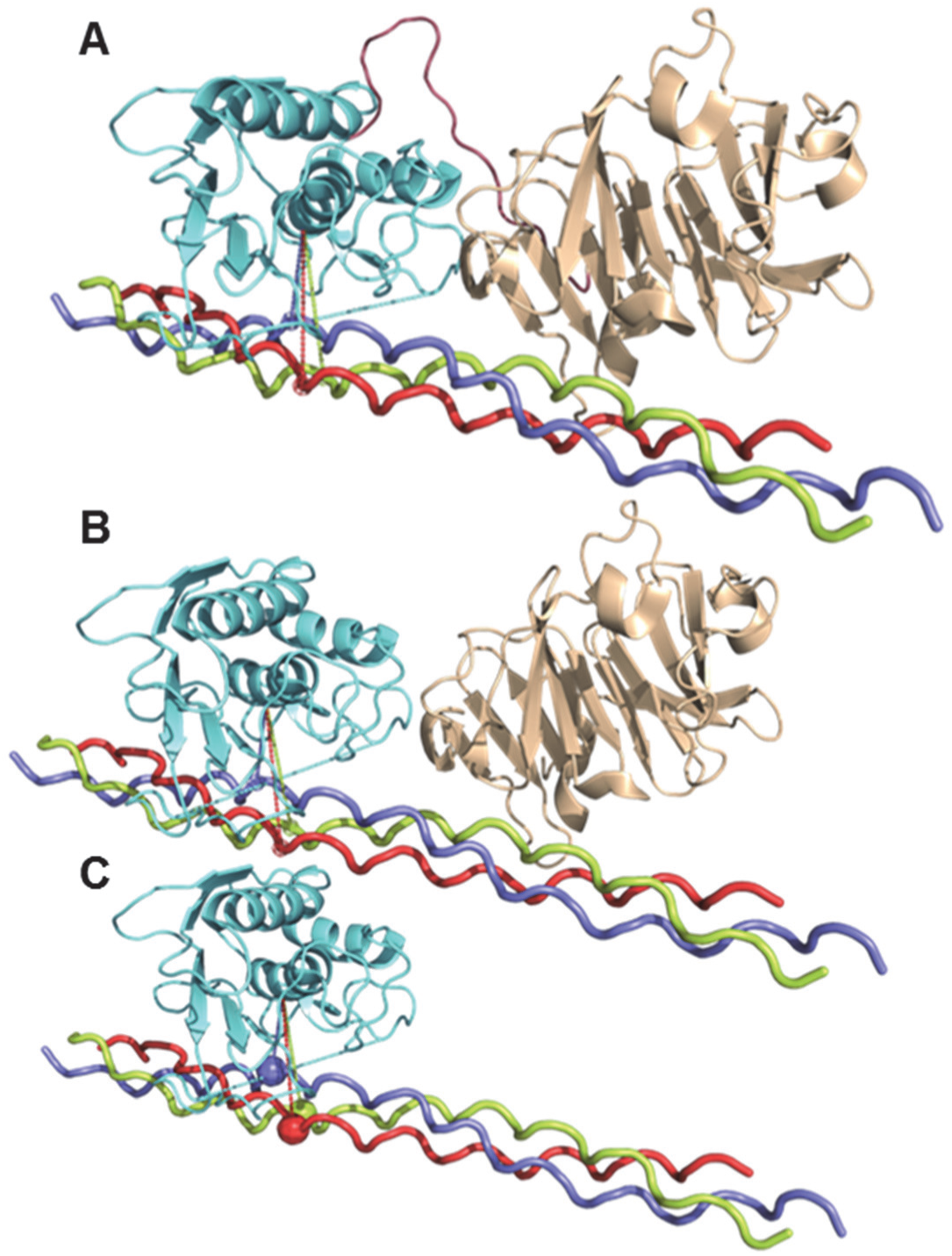
Inter-domain communications via collagen even without the linker. ANM simulations of the catalytic pocket opening as measured by the distance between N171 and T230 (cyan line) are (**A**) 2.56±0.52 nm, (**B**) 2.56±0.26 nm, and (**C**) 2.55±0.19 nm. The error bars represent the stds of 60 measurements of the catalytic pocket opening obtained from 20 frames each for the three slowest normal modes. The catalytic domain (cyan), the linker (brown), and the hemopexin (wheat) domains represent the residue ranges F100-Y260, G261-C278, and D279-C466, respectively.

In summary, we measured the inter-domain dynamics of MMP1 on type-1 collagen fibrils at the single-molecule level. To distinguish the functionally relevant MMP1 conformations, we used inactive MMP1, an enhancer of MMP1 activity (MMP9), and an inhibitor of MMP1 activity (tetracycline). We found that an open conformation, which has the two domains well-separated, is essential for function. Tetracycline forms two hydrogen bonds with both domains and inhibits dynamics resulting in similar histograms for active and inactive MMP1. Two-state stochastic simulations, coarse-grained ANM simulations, and all-atom MD simulations reproduced some features observed in single-molecule measurements. We modeled MMP1 dynamics as a two-state stochastic system, which enabled using histograms and autocorrelations to calculate the kinetic rates. MD and ANM simulations also provided new insights into MMP1 dynamics to show that a larger inter-domain motion of MMP1 correlates with a larger catalytic pocket opening of the catalytic domain and a smaller rms distance between the MMP1 catalytic site and the cleavage sites on collagen.

Our results resolve the debate whether the two MMP1 domains are well-separated (open low FRET state) or not (closed high FRET state). Based on NMR and SAXS experiments and the all-atom MD simulation studies (13, 15-17), previous studies argued that a larger separation between the catalytic and hemopexin domains is necessary for activity. In contrast, X-ray crystallography suggested that the two domains need to be closer (12). Also, it is well-established that MMP1 exists in an equilibrium between open and closed conformations (15, 16, 82). Our single molecule measurements, all-atom simulations, and two-state stochastic simulations also support that MMP1 exists in equilibrium between the two conformations. However, the ratio between the two states differs between active and inactive MMP1. Our results suggest that active MMP1 prefers the open conformation, whereas inactive MMP1 prefers the closed state. Since the crystal structure of MMP1 bound to a model of triple-helical collagen (4AUO) used inactive MMP1, it is not surprising that X-ray crystallography suggests a preference for the closed conformation (12).

In contrast to previous studies on water-soluble triple-helical collagen monomers, we measured MMP1 dynamics on water-insoluble type-1 collagen fibrils using smFRET. MMP1 dynamics on fibrils show similarities with dynamics on monomers and support previous mechanistic insights obtained from monomer-based studies. However, we know that MMP1 activity on monomers and fibrils are different. To this end, we assumed that monomers in solution will be more flexible than monomers in fibrils and performed MD simulations with restrained and unrestrained collagen backbones. Interestingly, inactive MMP1 did not show a significant change in preferred conformations between restrained and unrestrained collagen, but active MMP1 showed more preference for open conformations with the collagen backbone restrained. We found that two MMP1 domains can allosterically communicate to induce a larger catalytic pocket opening via the linker as well as the collagen. The E219Q point mutant in the catalytic domain significantly changed the fluctuations of residues in the hemopexin domain. Our approach is readily applicable to other multidomain proteins, including surface receptors, enzymes, and intracellular signaling proteins.

## Acknowledgments

The authors acknowledge John Czerski and Danielle Forristall for participating in the initial stage of smFRET measurements and analyses. Two grants to S.K.S. from the National Institutes of Health (RGM137295A) and the Colorado Office of Economic Development and International Trade (CTGG1 19-3508) partially supported this work.

## Author contributions

S.K.S. conceived and designed the overall project. L.K. and S.K.S. designed experiments. L.K. performed experiments. A.N. performed all-atom simulations. C.H. wrote codes for analyzing smFRET data and drew the TIRF schematics. J.P. performed ANM simulations and molecular docking. L.K., A.N., J.P., D.W., J.K.S., and S.K.S. analyzed data. S.K.S. wrote the manuscript. All authors read and edited the manuscript.

## Supplementary Information

**FIGURE S1.**
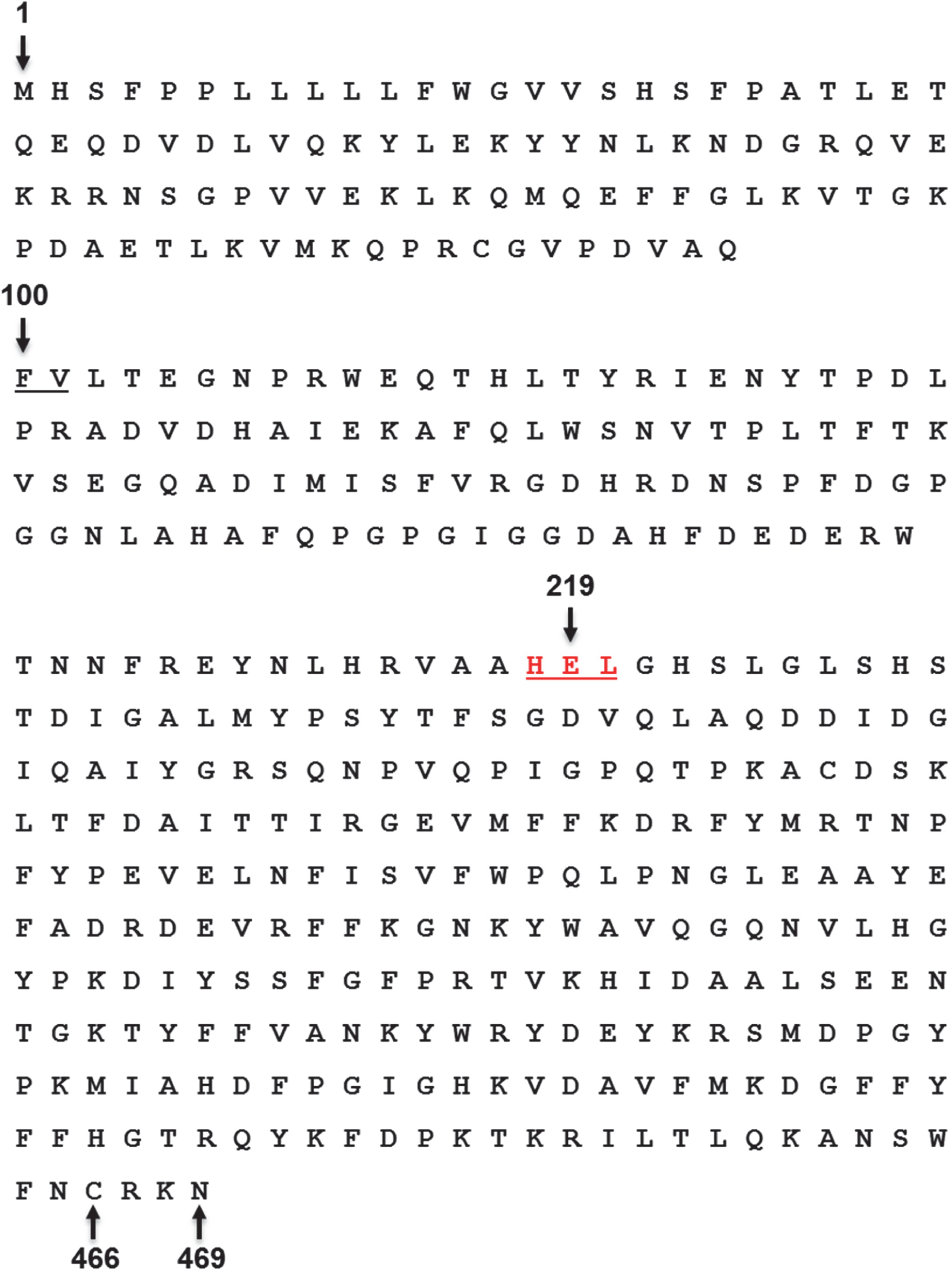
MMP1 sequence for experiments and simulations. Full-length MMP1 with the pro domain has 469 residues (M1-N469). Trypsin activates MMP1 by cleaving the F-V bond. Active and inactive MMP1 used in single molecule experiments has the same sequence (V101-N469) except the residue E219 at the catalytic site. For inactive MMP1, there is a single point mutation E219Q. For simulations, we used PDB ID 4AUO with the MMP1 sequence between F100 and C466. S142C, S366C, and E219Q mutations were modified in 4AUO to simulate the dynamics of active and inactive MMP1.

### AFM and SEM images of MMP-treated fibrils

We made reconstituted fibrils using RatCol® – Rat Tail Type-1 Collagen from Advanced Biomatrix by neutralizing the solution of type-1 collagen monomers and incubating at 37°C according to the manufacturer’s protocol.

**FIGURE S2.**
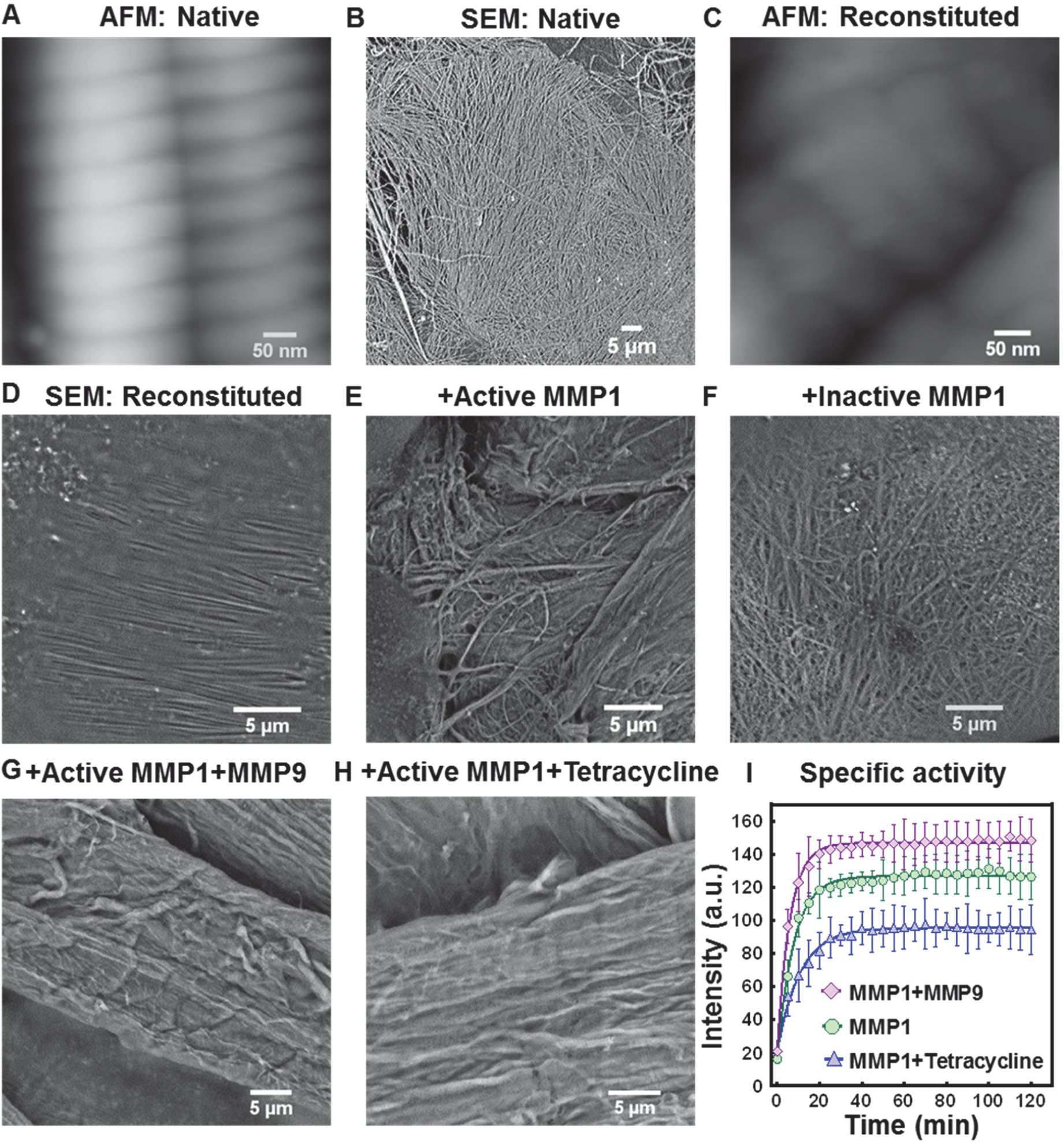
Activity of MMP1. (**A-H**) Surface morphology of MMP1-treated type-1 collagen fibrils. **(I)** Fluorescence from degraded peptide substrate, MCA-Lys-Pro-Leu-Gly-Leu-DPA-Ala-Arg-NH_2_, as a function of time for MMP1 (green circle), MMP1+MMP9 (magenta diamond), and MMP1+tetracycline (blue triangle). Solid lines are respective best fits to y=a-b*exp(-kt). The error bars are the standard deviations (std) of 3 technical repeats. After calibration, the specific activities are ∼1000 pmol/min/μg (MMP1), ∼1300 pmol/min/μg (MMP1+MMP9), ∼760 pmol/min/μg (MMP1+tetracycline).

We used native fibrils extracted from rat tails. We prepared three 200 μL reaction mixtures by adding 100 μL of 1 mg/mL active MMP1 with 1) 100 μL of protein buffer (50 mM Tris, 100 mM NaCl, pH 8.0), 2) 100 μL of 1 mg/mL MMP9, and 3) 100 μL of 100 μg/mL tetracycline. We prepared another 200 μL reaction mixture by adding 100 μL of 1 mg/mL inactive MMP1 with 100 μL of protein buffer. The slides were washed three times with 1 mL sterile distilled water to remove buffer precipitates. After washing, we air-dried the slides at room temperature. We incubated fibrils with these four reaction mixtures at 37° C for 4 h before imaging with AFM (Asylum MFP-3D Atomic Force Microscope) and SEM (Phenom Pro-Scanning Electron Microscope) (**Fig. S2, A-H**).

### Specific activity of MMP1 in the presence of MMP9 and tetracycline

We used a synthetic peptide substrate, MCA-Lys-Pro-Leu-Gly-Leu-DPA-Ala-Arg-NH_2_ (1), to measure the specific activity of MMP1 following the protocol described in a previous publication (2). MMP9 enhances the activity of MMP1, whereas tetracycline reduces the activity of MMP1 (**Fig. S2, I**).

### Best-fit parameters for experimental histograms and autocorrelations

We fitted two Gaussians to conformational histograms using the following equation:

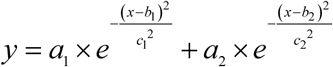

where a’s are amplitudes, b’s are centers, and c’s are widths of Gaussians. Best-fit parameters are in **Table S1**, with the centers highlighted in yellow. The error bars are the standard errors of mean (sem). The parameters b1 and b2 are states S1 and S2, respectively.

**Table S1.** Best-fit parameters for Gaussian fits to histograms in Fig. 2, A-C.

| MMP1 without ligands |  |  | MMP1 with MMP9 |  | MMP1 with tetracycline |  |
| --- | --- | --- | --- | --- | --- | --- |
|  | Active | Inactive | Active | Inactive | Active | Inactive |
| a1 | 2.95±0.08 | 6.06±0.02 | 4.97±0.54 | 2.59±0.09 | 6.48±0.04 | 0.38±0.03 |
| b1/S1 | 0.44±0.01 | 0.56±0.01 | 0.43±0.01 | 0.52±0.01 | 0.51±0.01 | 0.42±0.01 |
| c1 | 0.08±0.01 | 0.09±0.01 | 0.08±0.01 | 0.10±0.01 | 0.08±0.01 | 0.05±0.01 |
| a2 | 3.50±0.06 | 0.25±0.05 | 2.91±0.74 | 5.75±0.08 | 0.39±0.04 | 5.48±0.01 |
| b2/S2 | 0.55±0.01 | 0.68±0.01 | 0.48±0.01 | 0.54±0.01 | 0.55±0.01 | 0.50±0.01 |
| c2 | 0.09±0.01 | 0.06±0.01 | 0.06±0.01 | 0.05±0.01 | 0.15±0.01 | 0.10±0.01 |

We calculated the autocorrelations of conformational fluctuations by using the following equation:

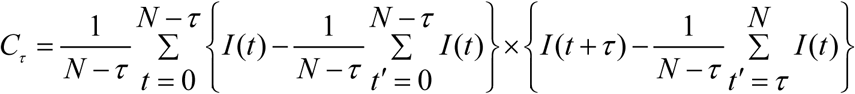

where *C*_*τ*_ is the autocorrelation at lag number *τ, N* is the total number of data points in a FRET trajectory, and *I* (*t*) is the FRET value at time point *t*.

We normalized autocorrelations by dividing autocorrelations at each lag by *C*_*τ* =0_. We fitted autocorrelations between *τ* = 1 and *τ* = 1000 to both power law and exponential. We used a form of Pareto distribution (3) as power law that satisfies the boundary conditions of our calculated autocorrelations, i.e., *C*_*τ* = 0_ = 1at *t* = 0 and *C*_*τ* = ∞_ = 0 at *t* = ∞. The power law and exponential functions used are:

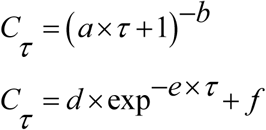

Generally, the power law does not fit the autocorrelations of trajectories obtained from experiments and two-state Poisson process simulations. Nevertheless, we have included them for comparison because power law fits better with dynamics simulated using all-atom MD (**Fig. 3**).

**Table S2.** Best-fit parameters for exponential fits to autocorrelations in Fig. 2, D-F.

| MMP1 without ligands |  |  | MMP1 with MMP9 |  | MMP1 with tetracycline |  |
| --- | --- | --- | --- | --- | --- | --- |
|  | Active | Inactive | Active | Inactive | Active | Inactive |
| d | 0.57±0.01 | 0.34±0.01 | 0.32±0.01 | 0.13±0.01 | 0.24±0.01 | 0.42±0.01 |
| e | 0.13±0.01 | 0.08±0.01 | 0.03±0.01 | 0.09±0.01 | 0.08±0.01 | 0.05±0.01 |
| f | -0.02±0.01 | -0.01±0.01 | 0.01±0.01 | 0.01±0.01 | -0.01±0.01 | -0.01±0.01 |

Best-fit parameters for exponential fits to experimental autocorrelations are in **Table S2**. The error bars are sems. The calculated rates from histograms and autocorrelations are in **Table S3**

**Table S3.**
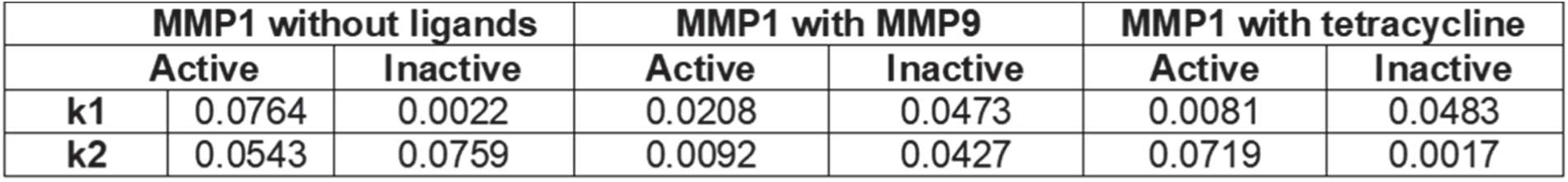
Calculated kinetic rates (s^−1^) of interconversion between states.

**FIGURE S3.**
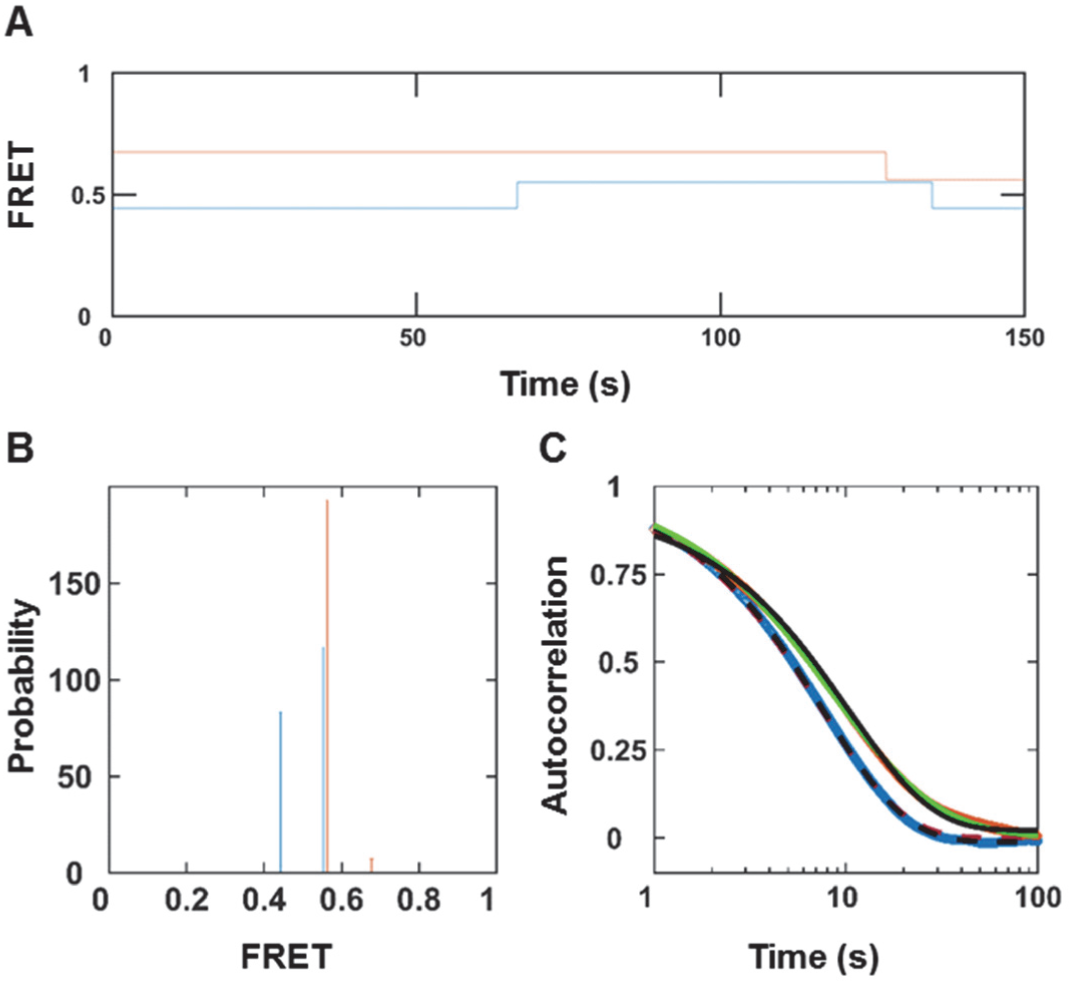
MMP1 inter-domain dynamics as a Poisson process without noise. (**A**) An example of simulated two-state FRET trajectory without noise for active (blue) and inactive (orange) MMP1. (**B**) Histograms of the recovered FRET values with bin size=0.005. (**C**) Autocorrelations of simulated trajectories recover the sum, k1+k2, from exponential fits (active: dashed black line; inactive: solid black line). As expected, power law does not fit autocorrelations (active: dashed red line; inactive: solid green line). The error bars are the sems for histograms and autocorrelations and are too small to be seen.

### Best-fit parameters for two-state Poisson process simulations

With and without noise, the peak positions (**Table S4**) determined from **Fig. 3, B** and **Fig. S3, B**, respectively, matched with the input FRET values. The exponential fits to autocorrelations recovered the expected sum of k1 and k2 (**Table S5**, right panel). Note that the power law reasonably fits the two-state Poisson process simulation without noise (**Fig. S3, C**). The error bars are the sems.

**Table S4.**
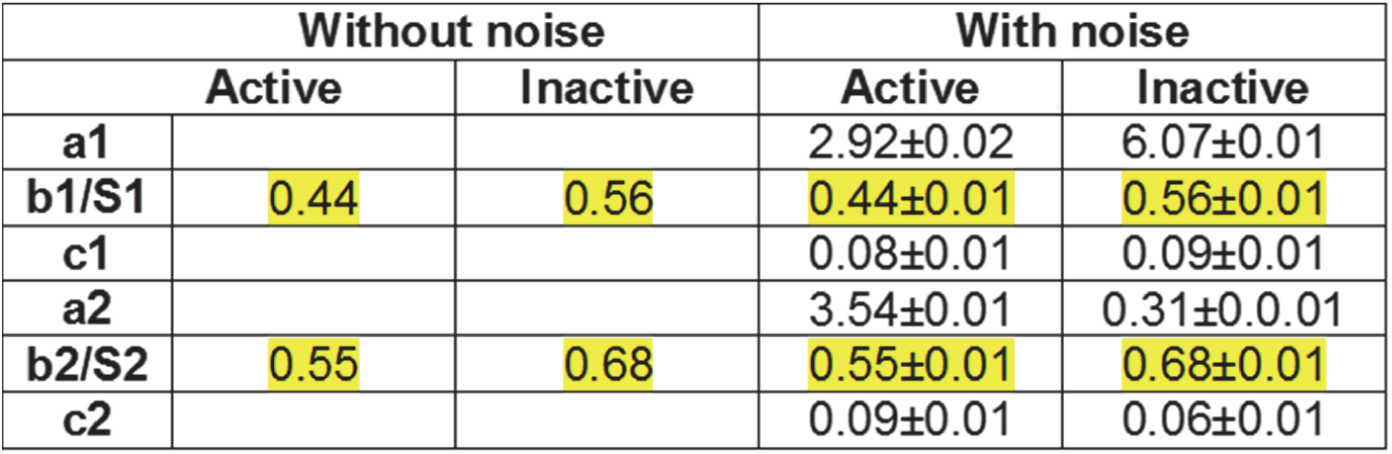
Gaussian fit parameters for two-state simulations in Fig. 3 and Fig. S3.

With noise, the peak positions and autocorrelation decay rates recovered the input parameters from Gaussian and exponential fits, respectively (**Table S4 and Table S5**). The error bars are the sems.

**Table S5.**
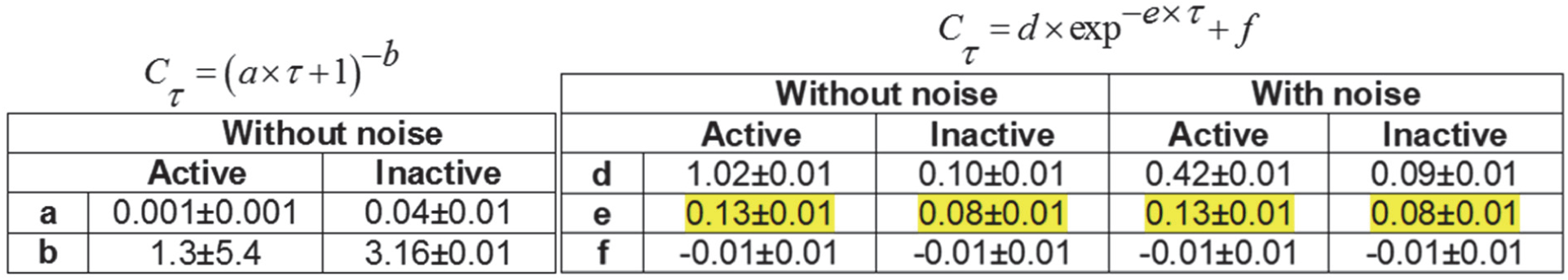
Fit parameters for two-state simulation autocorrelations in Fig. 3 and Fig. S3.

### Best-fit parameters for all-atom MD simulations

For all-atom MD simulations, we calculated the distance between S142C and S366C in nm (**Table S6**). The error bars are the sems.

**Table S6.**
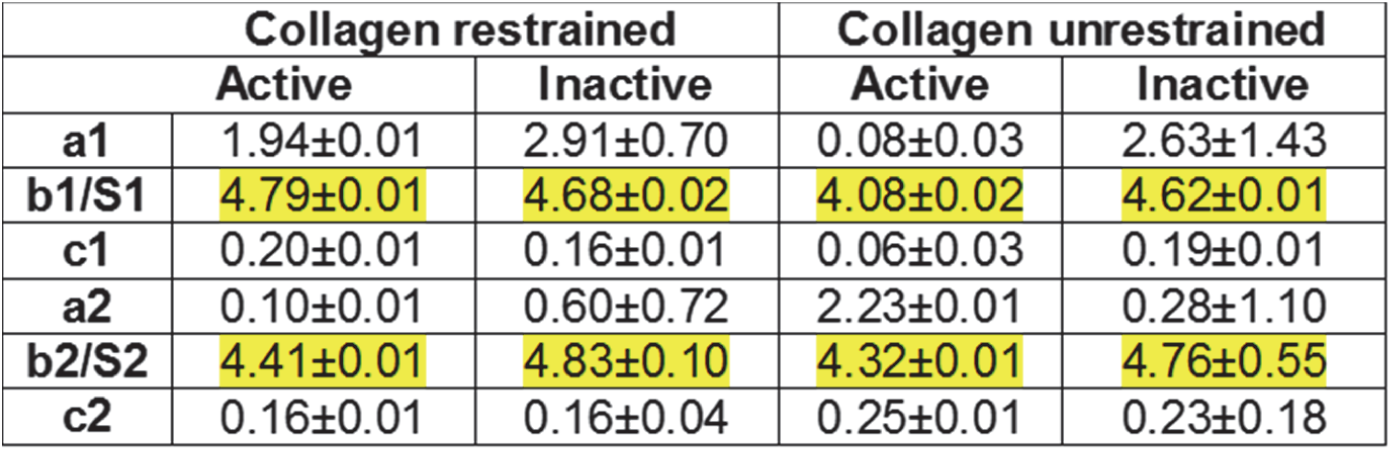
Best-fit parameters for Gaussian fits to MD histograms in Fig. 4.

**Table S7.**
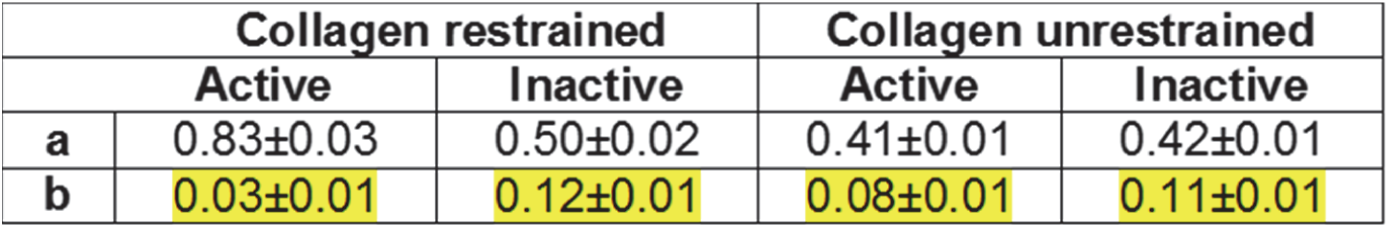
Best-fit parameters for power law fits to MD autocorrelations in Fig. 4.

### Molecular docking of tetracycline with MMP1

We docked tetracycline (ligand) with MMP1 (receptor, 4AUO) using Autodock Vina. We used the crystal structure 4AUO of MMP1 for docking. First, we removed the triple-helical collagen to expose the binding sites of MMP1 and selected a grid box by specifying the number of points and spacing in x, y, and z directions. The number of points in x, y, and z dimensions in MMP1 was 36, 52, and 54. Also, the spacings (Angstrom) in x, y, and z directions were 16.8, 120.1, and 23.5. The docking poses revealed two clusters of docking: one at the active site and the other between the catalytic and hemopexin domains (**Fig. S4, A**). Tetracycline formed two hydrogen bonds with PHE207 and GLU209 in the catalytic domain, and two hydrogen bonds with VAL319 and GLN323 in the hemopexin domain (**Fig. S4, B**). The presence of these hydrogen bonds suggests that tetracycline may restrict inter-domain motion of both active and inactive MMP1. The presence of these hydrogen bonds may explain the similar conformational histograms observed in experiments (**Fig. 2, A-C**).

**FIGURE S4.**
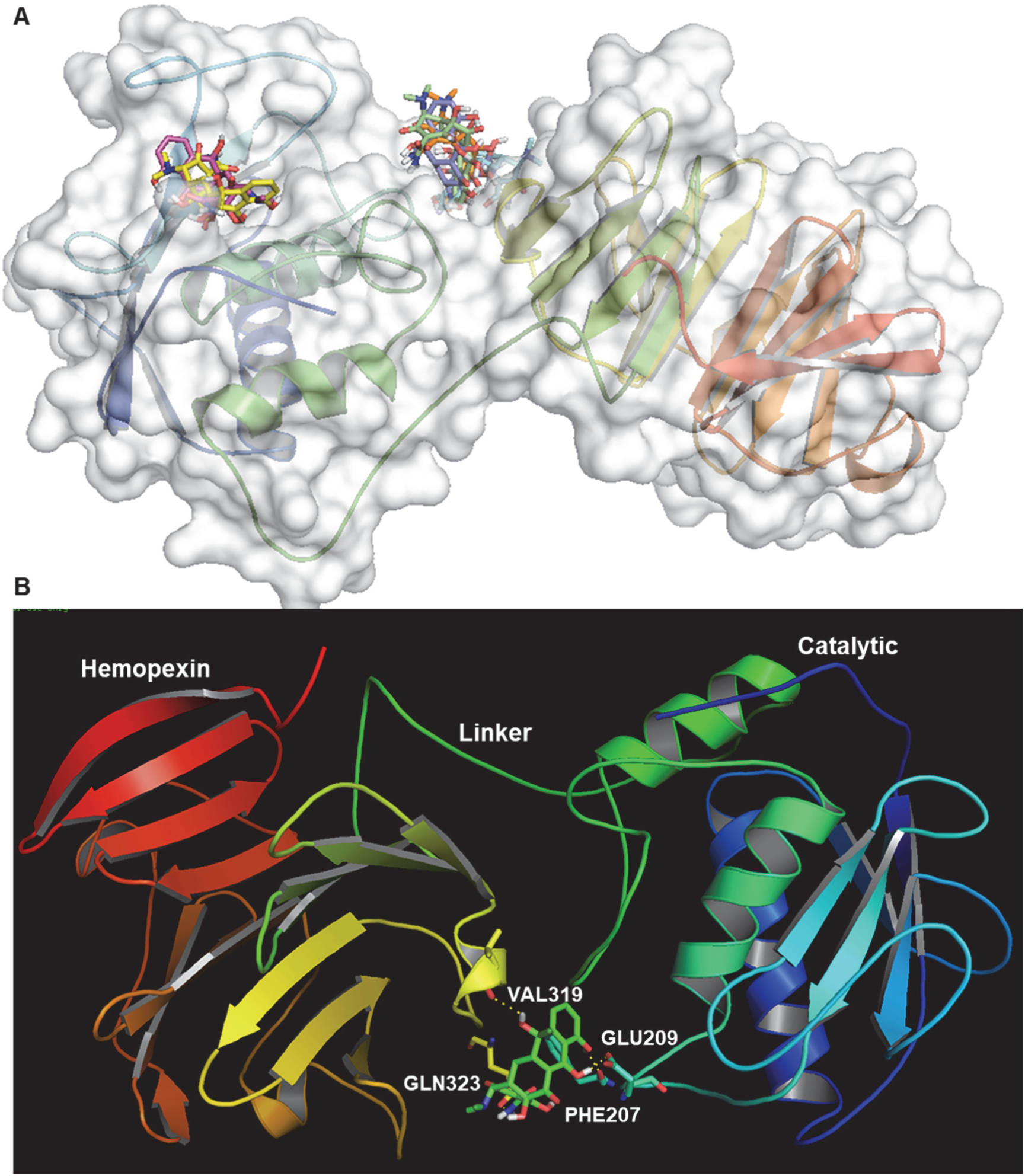
Molecular docking of tetracycline with MMP1. (**A**) Two clusters of docking poses near the catalytic site and linker region. (**B**) The docking pose with the highest docking score of −7.5 kcal/mol. Tetracycline forms two hydrogen bonds with both the catalytic and hemopexin domains, which may restrict inter-domain dynamics.

### Root-mean-squared fluctuation of amino acids from all-atom MD simulations

We quantified the conformational changes for active (E219) and inactive (E219Q) from simulations with the collagen backbone unrestrained. For both active and inactive MMP1, we modified 4AUO to include S142C and S366C mutations. Also, 4AUO has Q at 219, which we changed to E to simulate the effect of active MMP1. Note that 4AUO has amino acids between F100 and C466, which means 4AUO has the pro domains (amino acids between N1 and Q99) removed for both active and inactive MMP1. A sampling of simulated trajectories showed differences in the side-chain fluctuations and conformational space explorations. To quantify the difference in variations, we calculated the root-mean-squared fluctuation (RMSF) for active and inactive MMP1.

**FIGURE S5.**
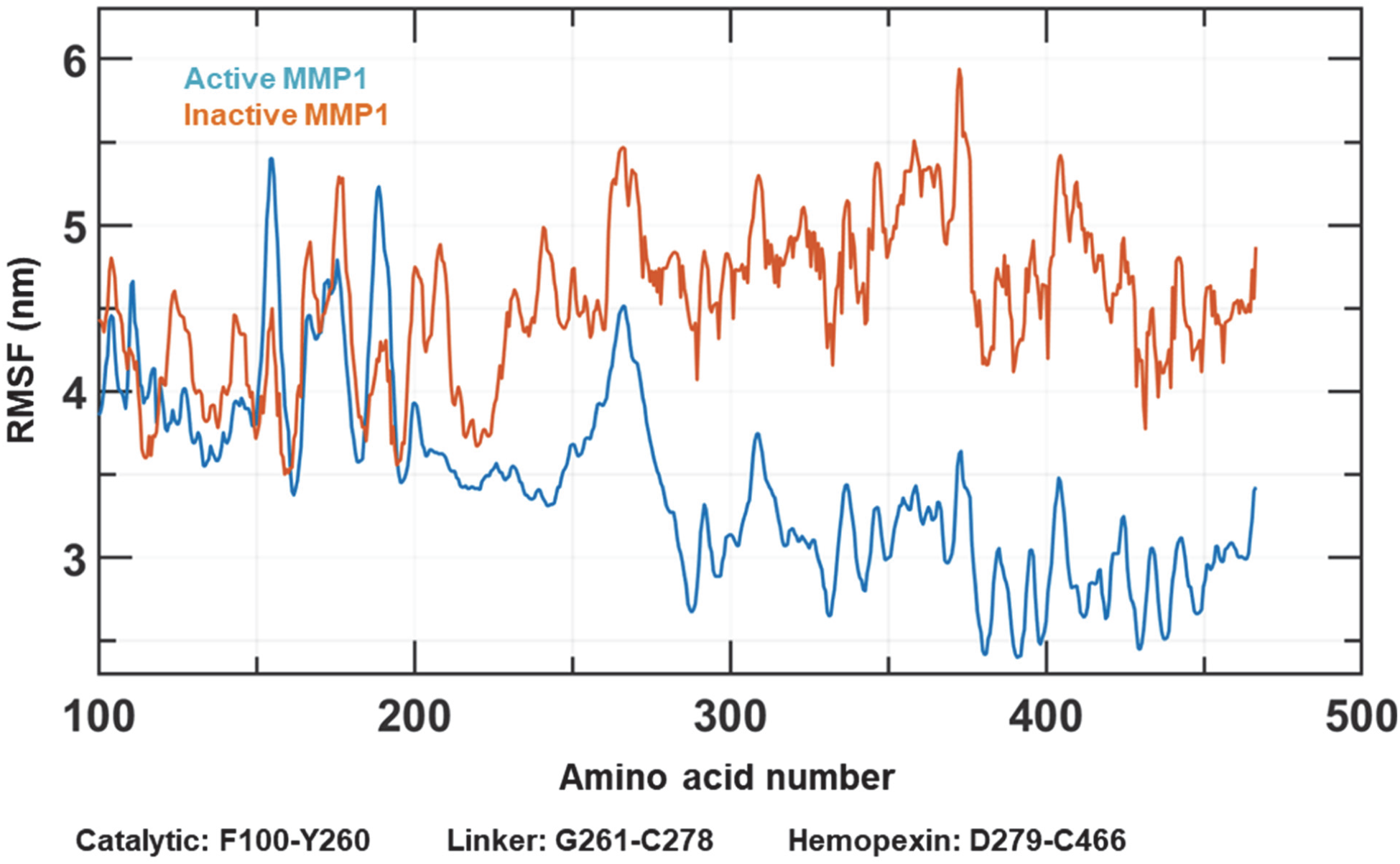
Analysis of fluctuations. The RMSF of amino acids for active (blue) and inactive (orange) MMP1 measured with respect to the energy-minimized structure before any simulation integration steps. During the 700 ns-long simulations with time step 2 fs, we saved data every 5 ps with the collagen unrestrained. The catalytic domain, the linker, and the hemopexin domains are defined roughly in the ranges F100-Y260, G261-C278, and D279-C466, respectively. The means±std deviations of RMSF across all the amino acids are 3.6±0.7 nm for active and 4.6±0.4 nm for catalytically inactive MMP1.

We calculated RMSF of MMP1 from the NPT simulation with the collagen backbone unrestrained using the GROMACS gmx toolset. We used the initial energy-minimized conformation before MD integration as a reference structure. In both active and inactive simulations, the catalytic domains showed a similar degree of side-chain fluctuation. However, the linker and hemopexin domains showed significant differences in the side-chain fluctuations. The changes in the hemopexin domain due to the mutation E219Q in the catalytic domain suggest communications between MMP1 domains. Such allosteric communications are in contrast to a previous NMR-based report on the effect of E219A on MMP12, another of the MMP family, interacting with type-V collagen that suggested no influence outside the catalytic cleft (4). In other words, allosteric communications depend on MMPs and the type of collagen. Further, the higher std of fluctuations across all amino acids for active MMP1 in comparison to inactive MMP1 is consistent with the wider experimental histograms for active MMP1 (**Fig. 2, A-C**). Surprisingly, allosteric communications appear within 700 ns of simulations shown in **Fig. S5**. However, previous studies have shown that single point mutations can result in protein sampling through different regions of the conformational phase space in short simulation times (5). The difference in RMSF may arise because the size differences between the amino acid substitutions may result in a change in van der Waals interaction (6). The differences could also be the result of a change in hydrogen bond configuration, as reported, for example, on the effects of active site mutations in cyp51A protein leading to resistance against therapeutics (7). Further studies are needed to define the mechanism behind the observed RMSF differences.

### Principal Component Analysis

Protein dynamics involve a wide range of length and time scales. For example, our MD simulations calculate Angstrom-scale motion with 2 fs time resolution over durations of a few hundred nanoseconds. In contrast, our smFRET measurements determine nanometer-scale movement with 100 ms time resolution over durations of a few hundred seconds. One approach to reveal the connections in such a wide range of scales is to note that only a few modes out of many protein conformations contain the majority of fluctuations. To find these critical (slow) modes, principal component analysis (PCA) is one of the most efficient methods. PCA involves the calculation of a covariance matrix followed by eigenvalue calculation of the covariance matrix to reduce a multidimensional complex set of variables to a few principal components (8).

**FIGURE S6.**
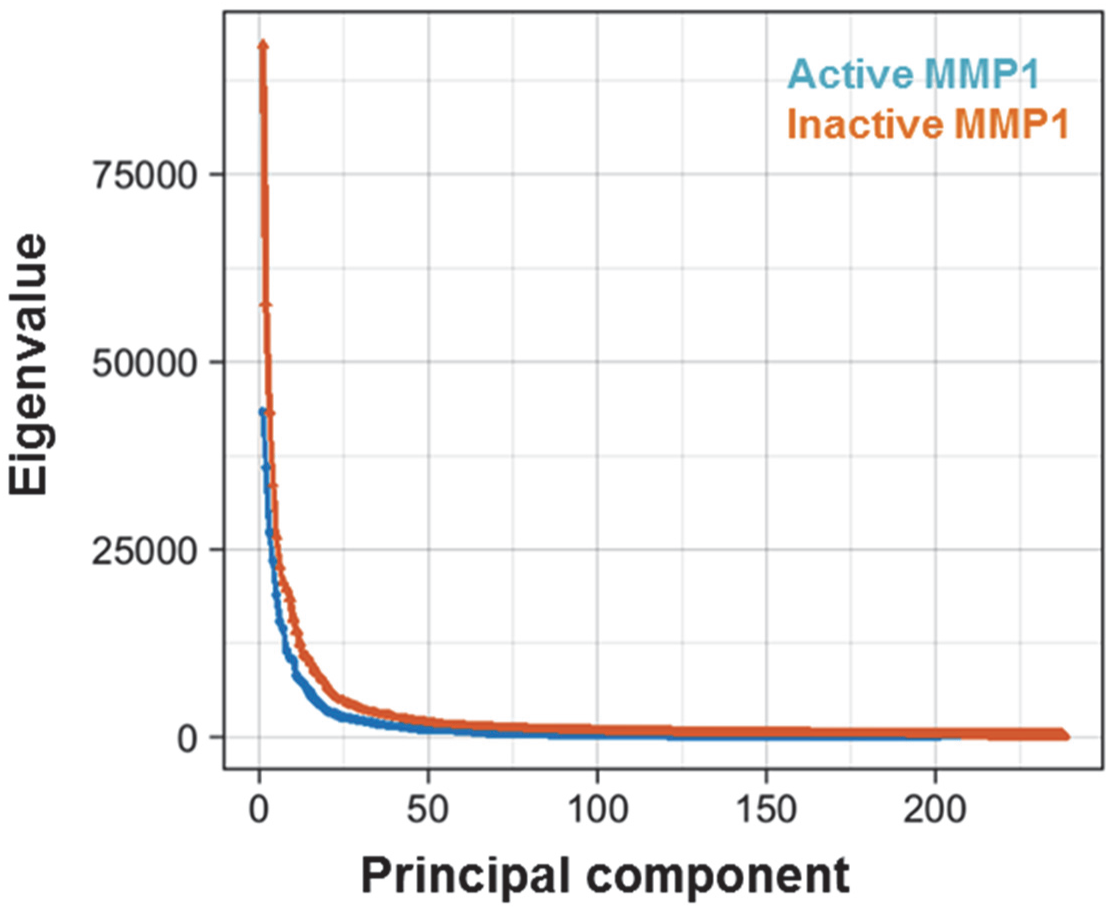
Principal component analysis. The distribution of eigenvalues as a function of principal components for active and inactive MMP1. The highest variations in protein dynamics occur in the first twenty principal components.

We performed protein principal component analysis to capture the overall dynamics of active and inactive MMP1 with the collagen unrestrained. We used the alpha-carbons (peptide-backbone carbon adjoining each side chain) in the 700 ns-long simulations. We used the pyPcazip software (9) to compress each trajectory file (0.9 of the original file) and extracted PCA-related metrics using pyPczdump. We generated simulation files (.xtc) without water and removed collagen, ions, and periodic boundary artifacts by treating each simulation with the gmx trjconv GROMACS utility. The majority of fluctuations in MMP1 dynamics falls within the first twenty principal components (**Fig. S6**), which is typical for proteins in general (10).

We analyzed the first four PCAs for active (**Fig. S7**) and inactive MMP1 (**Fig. S8**). Although MMP1 samples conformations around the initial structure, the first four principal components for active MMP1 revealed only one primary populated state (**Fig. S7**). In comparison, the PCA analysis for inactive MMP1 revealed a second populated state (**Fig. S8**), different from the conformation closer to the initial structure. The deviation between the primary populated states compares well with the difference in side-chain fluctuation between active and inactive MMP1.

**FIGURE S7.**
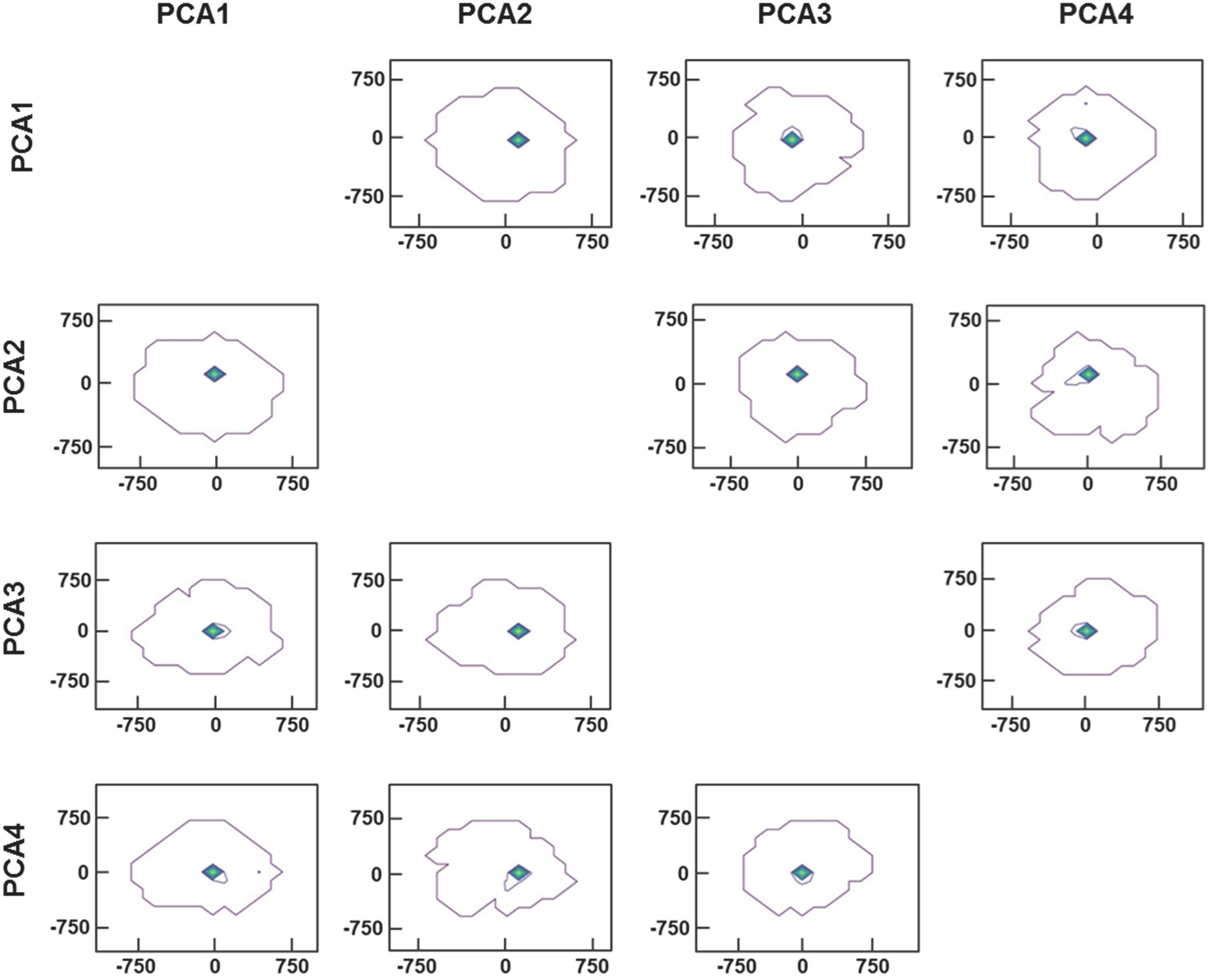
Principal component analysis for active MMP1. Conformational sampling plots for active MMP1 projected onto the first four principal components. One primary populated state can be distinguished.

**FIGURE S8.**
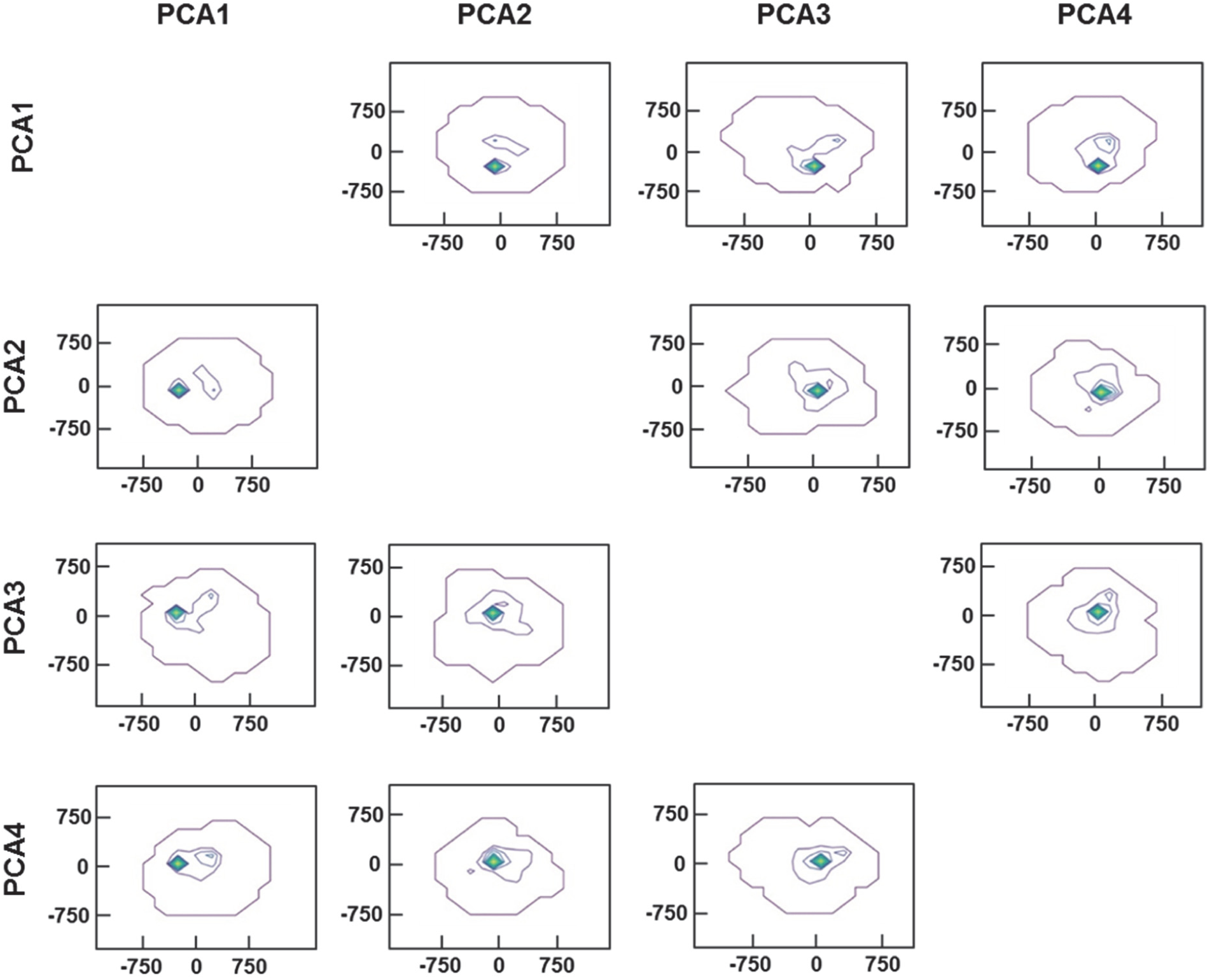
Principal component analysis for inactive MMP1. Conformational sampling plots for inactive MMP1 projected onto the first four principal components. The primary populated state accompanies a second less populated state.

## References

1. Karplus, M. 2010. Role of conformation transitions in adenylate kinase. Proceedings of the National Academy of Sciences 107(17):E71–E71.

2. Pisliakov, A. V., J. Cao, S. C. Kamerlin, and A. Warshel. 2009. Enzyme millisecond conformational dynamics do not catalyze the chemical step. Proceedings of the National Academy of Sciences 106(41):17359–17364.

3. Henzler-Wildman, K. A., M. Lei, V. Thai, S. J. Kerns, M. Karplus, and D. Kern. 2007. A hierarchy of timescales in protein dynamics is linked to enzyme catalysis. Nature 450(7171):913.

4. Chung, L. D., D. Dinakarpandian, N. Yoshida, J. L. Lauer-Fields, G. B. Fields, R. Visse, and H. Nagase. 2004. Collagenase unwinds triple-helical collagen prior to peptide bond hydrolysis. Embo Journal 23(15):3020–3030.

5. Page-McCaw, A., A. J. Ewald, and Z. Werb. 2007. Matrix metalloproteinases and the regulation of tissue remodelling. Nature reviews. Molecular cell biology 8(3):221–233.

6. Jackson, B. C., D. W. Nebert, and V. Vasiliou. 2010. Update of human and mouse matrix metalloproteinase families. Human genomics 4(3):194.

7. Ratnikov, B. I., P. Cieplak, K. Gramatikoff, J. Pierce, A. Eroshkin, Y. Igarashi, M. Kazanov, Q. Sun, A. Godzik, and A. Osterman. 2014. Basis for substrate recognition and distinction by matrix metalloproteinases. Proceedings of the National Academy of Sciences 111(40):E4148–E4155.

8. Gunasekaran, K., B. Ma, and R. Nussinov. 2004. Is allostery an intrinsic property of all dynamic proteins? Proteins: Structure, Function, and Bioinformatics 57(3):433–443.

9. Kadler, K. E. 2017. Fell Muir Review: Collagen fibril formation in vitro and in vivo. International journal of experimental pathology.

10. Nerenberg, P. S., R. Salsas-Escat, and C. M. Stultz. 2008. Do collagenases unwind triple-helical collagen before peptide bond hydrolysis? Reinterpreting experimental observations with mathematical models. Proteins: Structure, Function, and Bioinformatics 70(4):1154–1161.

11. Leikina, E., M. V. Mertts, N. Kuznetsova, and S. Leikin. 2002. Type I collagen is thermally unstable at body temperature. Proceedings of the National Academy of Sciences of the United States of America 99(3):1314–1318.

12. Manka, S. W., F. Carafoli, R. Visse, D. Bihan, N. Raynal, R. W. Farndale, G. Murphy, J. J. Enghild, E. Hohenester, and H. Nagase. 2012. Structural insights into triple-helical collagen cleavage by matrix metalloproteinase 1. Proc Natl Acad Sci U S A 109(31):12461–12466.

13. Karabencheva-Christova, T. G., C. Z. Christov, and G. B. Fields. 2017. Conformational Dynamics of Matrix Metalloproteinase-1· Triple-Helical Peptide Complexes. The Journal of Physical Chemistry B.

14. Manka, S. W., D. Bihan, and R. W. Farndale. 2019. Structural studies of the MMP-3 interaction with triple-helical collagen introduce new roles for the enzyme in tissue remodelling. Scientific Reports 9(1):1–14.

15. Bertini, I., M. Fragai, C. Luchinat, M. Melikian, M. Toccafondi, J. L. Lauer, and G. B. Fields. 2012. Structural basis for matrix metalloproteinase 1-catalyzed collagenolysis. Journal of the American Chemical Society 134(4):2100–2110.

16. Cerofolini, L., G. B. Fields, M. Fragai, C. F. Geraldes, C. Luchinat, G. Parigi, E. Ravera, D. I. Svergun, and J. M. Teixeira. 2013. Examination of matrix Metalloproteinase-1 in solution a preference for the pre-collagenolysis state. Journal of Biological Chemistry 288(42):30659–30671.

17. Arnold, L. H., L. E. Butt, S. H. Prior, C. M. Read, G. B. Fields, and A. R. Pickford. 2011. The interface between catalytic and hemopexin domains in matrix metalloproteinase-1 conceals a collagen binding exosite. Journal of Biological Chemistry 286(52):45073–45082.

18. Dittmore, A., J. Silver, S. K. Sarkar, B. Marmer, G. I. Goldberg, and K. C. Neuman. 2016. Internal strain drives spontaneous periodic buckling in collagen and regulates remodeling. Proceedings of the National Academy of Sciences:201523228.

19. Sarkar, S. K., B. Marmer, G. Goldberg, and K. C. Neuman. 2012. Single-molecule tracking of collagenase on native type I collagen fibrils reveals degradation mechanism. Current biology : CB 22(12):1047–1056.

20. Perumal, S., O. Antipova, and J. P. R. O. Orgel. 2008. Collagen fibril architecture, domain organization, and triple-helical conformation govern its proteolysis. Proceedings of the National Academy of Sciences of the United States of America 105(8):2824–2829.

21. Saffarian, S., I. E. Collier, B. L. Marmer, E. L. Elson, and G. Goldberg. 2004. Interstitial collagenase is a Brownian ratchet driven by proteolysis of collagen. Science 306(5693):108–111.

22. Greenwald, R., L. Golub, N. Ramamurthy, M. Chowdhury, S. Moak, and T. Sorsa. 1998. In vitro sensitivity of the three mammalian collagenases to tetracycline inhibition: relationship to bone and cartilage degradation. Bone 22(1):33–38.

23. Golub, L., N. Ramamurthy, T. McNamara, B. Gomes, M. Wolff, A. Casino, A. Kapoor, J. Zambon, S. Ciancio, and M. Schneir. 1984. Tetracyclines inhibit tissue collagenase activity. Journal of periodontal research 19(6):651–655.

24. Goldberg, G. I., A. Strongin, I. Collier, L. Genrich, and B. Marmer. 1992. Interaction of 92-kDa type IV collagenase with the tissue inhibitor of metalloproteinases prevents dimerization, complex formation with interstitial collagenase, and activation of the proenzyme with stromelysin. Journal of Biological Chemistry 267(7):4583–4591.

25. Messens, J., and J.-F. Collet. 2006. Pathways of disulfide bond formation in Escherichia coli. The international journal of biochemistry & cell biology 38(7):1050–1062.

26. Rowsell, S., P. Hawtin, C. A. Minshull, H. Jepson, S. M. Brockbank, D. G. Barratt, A. M. Slater, W. L. McPheat, D. Waterson, and A. M. Henney. 2002. Crystal structure of human MMP9 in complex with a reverse hydroxamate inhibitor. Journal of molecular biology 319(1):173–181.

27. Kumar, L., W. Colomb, J. Czerski, C. R. Cox, and S. K. Sarkar. 2018. Efficient protease based purification of recombinant matrix metalloprotease-1 in E. coli. Protein expression and purification 148:59–67.

28. Stennett, E. M., M. A. Ciuba, and M. Levitus. 2014. Photophysical processes in single molecule organic fluorescent probes. Chemical Society Reviews 43(4):1057–1075.

29. Cordes, T., J. Vogelsang, and P. Tinnefeld. 2009. On the mechanism of Trolox as antiblinking and antibleaching reagent. Journal of the American Chemical Society 131(14):5018–5019.

30. Rasnik, I., S. A. McKinney, and T. Ha. 2006. Nonblinking and long-lasting single-molecule fluorescence imaging. Nature methods 3(11):891.

31. Kochevar, I. E., and R. W. Redmond. 2000. Photosensitized production of singlet oxygen. Methods in enzymology. Elsevier, pp. 20–28.

32. Best, R. B., K. A. Merchant, I. V. Gopich, B. Schuler, A. Bax, and W. A. Eaton. 2007. Effect of flexibility and cis residues in single-molecule FRET studies of polyproline. Proceedings of the National Academy of Sciences 104(48):18964–18969.

33. Levitus, M., and S. Ranjit. 2011. Cyanine dyes in biophysical research: the photophysics of polymethine fluorescent dyes in biomolecular environments. Quarterly reviews of biophysics 44(1):123–151.

34. Merchant, K. A., R. B. Best, J. M. Louis, I. V. Gopich, and W. A. Eaton. 2007. Characterizing the unfolded states of proteins using single-molecule FRET spectroscopy and molecular simulations. Proceedings of the National Academy of Sciences 104(5):1528–1533.

35. Abraham, M. J., T. Murtola, R. Schulz, S. Páll, J. C. Smith, B. Hess, and E. Lindahl. 2015. GROMACS: High performance molecular simulations through multi-level parallelism from laptops to supercomputers. SoftwareX 1:19–25.

36. Hess, B., H. Bekker, H. J. Berendsen, and J. G. Fraaije. 1997. LINCS: a linear constraint solver for molecular simulations. Journal of computational chemistry 18(12):1463–1472.

37. Nash, A., H. L. Birch, and N. H. de Leeuw. 2017. Mapping intermolecular interactions and active site conformations: from human MMP-1 crystal structure to molecular dynamics free energy calculations. Journal of Biomolecular Structure and Dynamics 35(3):564–573.

38. Collier, T. A., A. Nash, H. L. Birch, and N. H. de Leeuw. 2016. Intra-molecular lysine-arginine derived advanced glycation end-product cross-linking in Type I collagen: A molecular dynamics simulation study. Biophysical chemistry 218:42–46.

39. Collier, T. A., A. Nash, H. L. Birch, and N. H. de Leeuw. 2015. Preferential sites for intramolecular glucosepane cross-link formation in type I collagen: A thermodynamic study. Matrix Biology 48:78–88.

40. Darden, T., D. York, and L. Pedersen. 1993. Particle mesh Ewald: An N· log (N) method for Ewald sums in large systems. The Journal of chemical physics 98(12):10089–10092.

41. Hoover, W. G. 1985. Canonical dynamics: Equilibrium phase-space distributions. Physical review A 31(3):1695.

42. Parrinello, M., and A. Rahman. 1981. Polymorphic transitions in single crystals: A new molecular dynamics method. Journal of Applied physics 52(12):7182–7190.

43. Eyal, E., L.-W. Yang, and I. Bahar. 2006. Anisotropic network model: systematic evaluation and a new web interface. Bioinformatics 22(21):2619–2627.

44. Isin, B., K. C. Tirupula, Z. N. Oltvai, J. Klein-Seetharaman, and I. Bahar. 2012. Identification of motions in membrane proteins by elastic network models and their experimental validation. Membrane Protein Structure and Dynamics. Springer, pp. 285–317.

45. Eyal, E., G. Lum, and I. Bahar. 2015. The anisotropic network model web server at 2015 (ANM 2.0). Bioinformatics 31(9):1487–1489.

46. Atilgan, A. R., S. Durell, R. L. Jernigan, M. Demirel, O. Keskin, and I. Bahar. 2001. Anisotropy of fluctuation dynamics of proteins with an elastic network model. Biophysical journal 80(1):505–515.

47. Haugland, R. P. 2002. Handbook of fluorescent probes and research products. Molecular Probes.

48. Neumann, U., H. Kubota, K. Frei, V. Ganu, and D. Leppert. 2004. Characterization of Mca-Lys-Pro-Leu-Gly-Leu-Dpa-Ala-Arg-NH2, a fluorogenic substrate with increased specificity constants for collagenases and tumor necrosis factor converting enzyme. Analytical biochemistry 328(2):166–173.

49. Kamerlin, S. C., and A. Warshel. 2010. Reply to Karplus: Conformational dynamics have no role in the chemical step. Proceedings of the National Academy of Sciences 107(17):E72–E72.

50. Patterson, M. L., S. J. Atkinson, V. Knäuper, and G. Murphy. 2001. Specific collagenolysis by gelatinase A, MMP-2, is determined by the hemopexin domain and not the fibronectin-like domain. FEBS letters 503(2-3):158–162.

51. Saito, S., M. J. Trovato, R. X. You, B. K. Lal, F. Fasehun, F. T. Padberg, R. W. Hobson, W. N. Duran, and P. J. Pappas. 2001. Role of matrix metalloproteinases 1, 2, and 9 and tissue inhibitor of matrix metalloproteinase-1 in chronic venous insufficiency. Journal of Vascular Surgery 34(5):930–937.

52. Asahi, M., K. Asahi, J. C. Jung, G. J. del Zoppo, M. E. Fini, and E. H. Lo. 2000. Role for matrix metalloproteinase 9 after focal cerebral ischemia, effects of gene knockout and enzyme inhibition with BB-94. Journal of Cerebral Blood Flow and Metabolism 20(12):1681–1689.

53. Fujimura, M., Y. Gasche, Y. Morita-Fujimura, J. Massengale, M. Kawase, and P. H. Chan. 1999. Early appearance of activated matrix metalloproteinase-9 and blood-brain barrier disruption in mice after focal cerebral ischemia and reperfusion. Brain Research 842(1):92–100.

54. Hanemaaijer, R., H. Visser, Y. T. Konttinen, P. Koolwijk, and J. H. Verheijen. 1998. A novel and simple immunocapture assay for determination of gelatinase-B (MMP-9) activities in biological fluids: Saliva from patients with Sjogren’s syndrome contain increased latent and active gelatinase-B levels. Matrix Biology 17(8-9):657–665.

55. Koshiba, T., R. Hosotani, M. Wada, K. Fujimoto, J. U. Lee, R. Doi, S. Arii, and M. Imamura. 1997. Detection of matrix metalloproteinase activity in human pancreatic cancer. Surgery Today-the Japanese Journal of Surgery 27(4):302–304.

56. Sakalihasan, N., P. Delvenne, B. V. Nusgens, R. Limet, and C. M. Lapiere. 1996. Activated forms of MMP(2) and MMP(9) in abdominal aortic aneurysms. Journal of Vascular Surgery 24(1):127–133.

57. Powell, B., D. C. Malaspina, I. Szleifer, and Y. Dhaher. 2019. Effect of collagenase– gelatinase ratio on the mechanical properties of a collagen fibril: a combined Monte Carlo– molecular dynamics study. Biomechanics and modeling in mechanobiology:1–11.

58. Jarymowycz, V. A., and M. J. Stone. 2006. Fast time scale dynamics of protein backbones: NMR relaxation methods, applications, and functional consequences. Chemical reviews 106(5):1624–1671.

59. Singh, W., G. B. Fields, C. Z. Christov, and T. G. Karabencheva-Christova. 2016. Importance of the Linker Region in Matrix Metalloproteinase-1 Domain Interactions. RSC advances 6(28):23223–23232.

60. Piccard, H., P. E. Van den Steen, and G. Opdenakker. 2007. Hemopexin domains as multifunctional liganding modules in matrix metalloproteinases and other proteins. Journal of leukocyte biology 81(4):870–892.

61. Overall, C. M. 2002. Molecular determinants of metalloproteinase substrate specificity. Molecular biotechnology 22(1):51–86.

62. Clark, I. M., and T. E. Cawston. 1989. Fragments of human fibroblast collagenase. Purification and characterization. Biochemical Journal 263(1):201–206.

63. Sarkar, S. K. 2016. Single Molecule Biophysics and Poisson Process Approach to Statistical Mechanics. Morgan & Claypool Publishers.

64. Zumbusch, A., L. Fleury, R. Brown, J. Bernard, and M. Orrit. 1993. Probing individual two-level systems in a polymer by correlation of single molecule fluorescence. Physical review letters 70(23):3584.

65. Wang, J., L. Xu, K. Xue, and E. Wang. 2008. Exploring the origin of power law distribution in single-molecule conformation dynamics: energy landscape perspectives. Chemical Physics Letters 463(4-6):405–409.

66. Tang, J., and R. Marcus. 2006. Chain dynamics and power-law distance fluctuations of single-molecule systems. Physical Review E 73(2):022102.

67. Maisuradze, G. G., A. Liwo, and H. A. Scheraga. 2009. Principal component analysis for protein folding dynamics. Journal of molecular biology 385(1):312–329.

68. Koshland Jr, D. E. 1995. The key–lock theory and the induced fit theory. Angewandte Chemie International Edition in English 33(23-24):2375–2378.

69. Koshland, D. E. 1958. Application of a theory of enzyme specificity to protein synthesis. Proceedings of the National Academy of Sciences 44(2):98–104.

70. Monod, J., J. Wyman, and J.-P. Changeux. 1965. On the nature of allosteric transitions: a plausible model. J Mol Biol 12(1):88–118.

71. Boehr, D. D., R. Nussinov, and P. E. Wright. 2009. The role of dynamic conformational ensembles in biomolecular recognition. Nature chemical biology 5(11):789–796.

72. Zwanzig, R. 1990. Rate processes with dynamical disorder. Accounts of Chemical Research 23(5):148–152.

73. Desiraju, G. R. 2002. Cryptic crystallography. Nature materials 1(2):77–79.

74. Stanley, H., S. Buldyrev, A. Goldberger, S. Havlin, C.-K. Peng, and M. Simons. 1993. Long-range power-law correlations in condensed matter physics and biophysics. Physica A: Statistical Mechanics and its Applications 200(1-4):4–24.

75. Buldyrev, S. V. 2006. Power law correlations in DNA sequences. Power laws, scale-free networks and genome biology. Springer, pp. 123–164.

76. Grau-Carles, P. 2001. Long-range power-law correlations in stock returns. Physica A: Statistical Mechanics and its Applications 299(3-4):521–527.

77. Stanley, H., S. Buldyrev, A. Goldberger, S. Havlin, C.-K. Peng, F. Sciortino, and M. Simons. 1993. Scaling concepts and complex fluids: long-range power-law correlations in DNA. Le Journal de Physique IV 3(C1):C1–15-C11-25.

78. Luscombe, N. M., J. Qian, Z. Zhang, T. Johnson, and M. Gerstein. 2002. The dominance of the population by a selected few: power-law behaviour applies to a wide variety of genomic properties. Genome biology 3(8):research0040. 0041.

79. Wertheim, G., M. Butler, K. West, and D. Buchanan. 1974. Determination of the Gaussian and Lorentzian content of experimental line shapes. Review of Scientific Instruments 45(11):1369–1371.

80. Colomb, W., J. Czerski, J. Sau, and S. Sarkar. 2017. Estimation of microscope drift using fluorescent nanodiamonds as fiducial markers. Journal of microscopy 266(3):298–306.

81. Chapeau-Blondeau, F. 2007. Autocorrelation versus entropy-based autoinformation for measuring dependence in random signal. Physica A: Statistical Mechanics and its Applications 380:1–18.

82. Bertini, I., M. Fragai, C. Luchinat, M. Melikian, E. Mylonas, N. Sarti, and D. I. Svergun. 2009. Interdomain flexibility in full-length matrix metalloproteinase-1 (MMP-1). Journal of Biological Chemistry 284(19):12821–12828.

## References

1. Neumann, U., H. Kubota, K. Frei, V. Ganu, and D. Leppert. 2004. Characterization of Mca-Lys-Pro-Leu-Gly-Leu-Dpa-Ala-Arg-NH2, a fluorogenic substrate with increased specificity constants for collagenases and tumor necrosis factor converting enzyme. Analytical biochemistry 328(2):166–173.

2. Kumar, L., W. Colomb, J. Czerski, C. R. Cox, and S. K. Sarkar. 2018. Efficient protease based purification of recombinant matrix metalloprotease-1 in E. coli. Protein expression and purification 148:59–67.

3. Arnold, B. C. 2014. Pareto distribution. Wiley StatsRef: Statistics Reference Online:1-10.

4. Prior, S. H., T. S. Byrne, D. Tokmina-Roszyk, G. B. Fields, and S. R. Van Doren. 2016. Path to Collagenolysis COLLAGEN V TRIPLE-HELIX MODEL BOUND PRODUCTIVELY AND IN ENCOUNTERS BY MATRIX METALLOPROTEINASE-Journal of Biological Chemistry 291(15):7888–7901.

5. Ng, H. W., C. A. Laughton, and S. W. Doughty. 2013. Molecular dynamics simulations of the adenosine A2a receptor: structural stability, sampling, and convergence. Journal of chemical information and modeling 53(5):1168–1178.

6. Loladze, V. V., D. N. Ermolenko, and G. I. Makhatadze. 2002. Thermodynamic consequences of burial of polar and non-polar amino acid residues in the protein interior. Journal of molecular biology 320(2):343–357.

7. Nash, A., and J. Rhodes. 2018. Simulations of CYP51A from Aspergillus fumigatus in a model bilayer provide insights into triazole drug resistance. Medical mycology 56(3):361–373.

8. Maisuradze, G. G., A. Liwo, and H. A. Scheraga. 2009. Principal component analysis for protein folding dynamics. Journal of molecular biology 385(1):312–329.

9. Shkurti, A., R. Goni, P. Andrio, E. Breitmoser, I. Bethune, M. Orozco, and C. A. Laughton. 2016. pyPcazip: A PCA-based toolkit for compression and analysis of molecular simulation data. SoftwareX 5:44–50.

10. David, C. C., and D. J. Jacobs. 2014. Principal component analysis: a method for determining the essential dynamics of proteins. Protein dynamics. Springer, pp. 193–226.

